# Stable epigenetic states set single-cell activation thresholds in mammalian expression systems

**DOI:** 10.64898/2026.09.04.749524

**Authors:** Eli J. Costa, Carolina Rios-Martinez, Cecelia J. Andrews, James E. Ferrell, Lacramioara Bintu

**Author notes:** These authors jointly supervised this work (James E. Ferrell, Lacramioara Bintu). Correspondence (J.E.F) and (L.B.).

## Abstract

Quantitatively relating transcription factor (TF) input to gene expression output is central to understanding mammalian gene regulation and essential for designing predictable synthetic expression systems. However, even minimal synthetic systems often exhibit unexplained behaviors. In a widely used inducible mammalian expression system, we show that transcriptional responses appear graded and sigmoidal at the population level but are largely all-or-none at the single-cell level. By combining single-cell sorting and single-molecule footprinting with mathematical modeling of transcriptional regulation, we found that this behavior is not caused by bursty transcription or bistability, but by long-lived, chromatin-encoded variability in TF occupancy and activation strength. This variability produced a range of activation thresholds in switch-like single-cell responses that were stable over time, resulting in bimodal gene expression across the population. These results advance our basic understanding of how TFs interact with chromatin to modulate quantitative features of single-cell and population level transcriptional responses.

## Introduction

Transcription factors (TFs) regulate gene expression programs by promoting or inhibiting the transcription of target genes, which then direct cell fates and functions^1,2^. The relationship between TF input (i.e. concentration) and gene expression output (i.e. mRNA levels) in part defines the regulatory capabilities of a given TF, and misregulation of gene expression is associated with a number of genetic diseases and disorders, including cancer^3,4^. Similarly, engineered TFs provide a powerful method for controlling cell behaviors in both basic research and therapeutic applications such as cell and gene therapies^5–7^. Developing a quantitative understanding of gene expression that is able to precisely relate TF input to transcriptional output in diverse contexts and applications remains a central goal in the field of gene regulation^8,9^.

TF-mediated gene regulation in mammals is a complex, multi-step process in which TFs recognize and bind to short DNA sequences within chromatin, which is itself a dynamic and epigenetically regulated structure. TFs then recruit various cofactors to mediate chromatin remodeling, RNA polymerase II binding, and transcription from gene promoters^1,3,10,11^. Because of this complexity and the number of distinct molecular mechanisms involved, obtaining a quantitative understanding of the relationship between TF input and gene expression output based on unifying fundamental principles has proven difficult^12^.

Synthetic expression systems are seemingly well suited for developing the foundation for a quantitative understanding of gene regulation and identifying these fundamental principles because, in some sense, they represent the simplest possible case of mammalian transcriptional control. Typically, in these systems, synthetic TFs (synTFs) orthogonal to the host genome bind proximally and specifically to cognate binding sites upstream of a minimal core promoter in an engineered target gene^5–7^. Many complex features of endogenous mammalian gene regulation, such as co-regulation by multiple TFs, distal enhancers, layered signaling, and feedback from gene regulatory networks are largely absent^13^. This simplicity, combined with a high level of control over the synTF and reporter gene, makes it possible to interrogate the effects of key variables on transcriptional responses in a controlled environment^8^. In addition, synthetic expression systems such as Tet-On are widely used as versatile and powerful tools in many areas of synthetic gene circuit engineering and fundamental molecular biology research (e.g. to tunably induce exogenous protein expression), making a quantitative understanding of their transcriptional response functions broadly relevant and valuable.

Despite their apparent simplicity, synthetic expression systems still exhibit unexplained behaviors and unexpected gene regulatory responses to synTF inputs. In particular, many groups have observed that transcriptional responses to increasing levels of a synTF are graded at the population level but largely all-or-none at the single-cell level^7,14–21^. This results in bimodal gene expression distributions, where increasing synTF induction primarily regulates the fraction of cells expressing a target gene rather than uniformly regulating the average expression level of all cells in the population (Figure 1A). Bimodality is observed for synthetic expression systems utilizing different DNA binding domains, transcriptional activation domains, and delivery/integration methods. It is unclear mechanistically what gives rise to this bimodality. Several potential mechanisms have been proposed, such as transcriptional bursting on slow timescales^12,16,22–25^ and chromatin-mediated bistability in single-cell transcriptional responses^26,27^; however, many quantitative models of gene regulation do not account for bimodality^28^ and in all cases the potential mechanisms remain to be rigorously tested in synthetic expression systems^8,9,29–36^.

**Figure 1:**
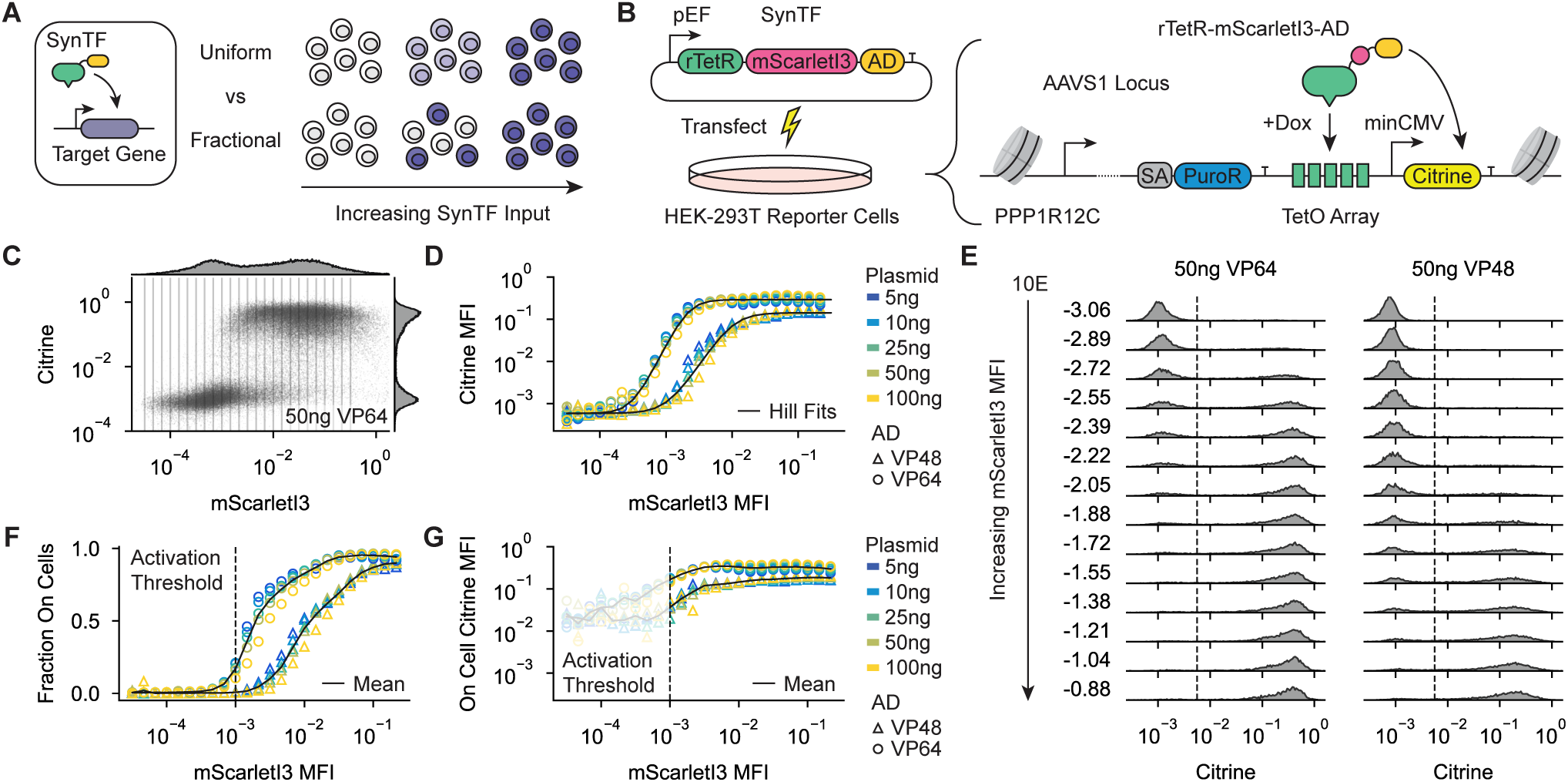
Transcriptional activation in the Tet-On system is graded at the population level but largely all-or-none at the single-cell level. A) Synthetic expression systems are often characterized by graded responses at the population level but fractional, all-or-none responses at the single-cell level. B) The Tet-On synthetic expression system used to quantify transcriptional response functions. The synTF consists of rTetR fused to mScarletI3 and an activation domain (AD, generally VP48 or VP64, as indicated). A Citrine reporter gene containing 9 TetO binding sites (unless otherwise indicated) upstream of a minimal promoter (minCMV) was integrated into the AAVS1 locus of HEK-293T cells and selected by promoter-trapping the upstream PPP1R12C promoter upon successful integration. SA = splice acceptor, PuroR = puromycin resistance marker. C) Example flow cytometry data obtained after transfecting 50ng of synTF (with the AD VP64) expression plasmid into reporter cells. 1000 ng/mL dox was added 24 hours after transfection, and measurements were taken 24 hours after dox addition. Fluorescence measurements were normalized by FSC and for bleed-through (Figure S1B). Vertical lines represent bins used to calculate the transcriptional response functions in D, F, and G. D) Average transcriptional response functions for transient transfections performed as in C. Citrine mean fluorescent intensity (MFI) was calculated across all cells falling within the mScarletI3 bins shown in C. mScarletI3 values are the mean of each bin. Solid lines are Hill function fits (Figure S1C). Shapes represent ADs; colors represent the amount of synTF expression plasmid transfected. Measurements between ADs were normalized to have the same background fluorescence (see Methods). E) Citrine expression distributions across mScarletI3 bins for transfections performed as in C. mScarletI3 values represent the mean of each bin. Dashed lines represent the threshold used to call active cells in F and G. F) Fraction of active cells within each mScarletI3 bin for transfections performed as in C. Solid lines are the mean fraction of active cells across transfections. Shapes represent ADs; colors represent the amount of synTF expression plasmid transfected. G) MFI of active cells within each mScarletI3 bin for transfections performed as in C. Solid lines are the mean MFI of active cells across transfections. Shapes represent ADs; colors represent the amount of synTF expression plasmid transfected. The shaded area denotes the region where very few cells in the population activate gene expression.

The inability to quantitatively describe gene expression responses even in simple, synthetic contexts indicates that our current understanding of mammalian gene regulation does not account for key features of transcriptional control. In addition, these shortcomings have practical implications for the design and use of synthetic gene circuits that commonly utilize these synthetic expression systems. Uncontrolled or heterogeneous expression can cause synthetic circuits to fail or confound the interpretation of experimental results^20,37–40^. Therefore, gaining a quantitative understanding of bimodality in synthetic expression systems would both advance our basic understanding of transcription in mammalian cells and address a major limitation in synthetic biology, improving therapeutic and research applications by enabling more predictable control over gene expression.

Here, we investigated single-cell and population-level transcriptional responses from the Tet-On synthetic expression system^41–43^ integrated at a safe-harbor locus in human cells. Consistent with previous reports, we observed switch-like activation at the single-cell level resulting in bimodal gene expression across the population. We found that bimodality was not caused by bistability or bursty transcription, but driven by cell-to-cell variability in ultrasensitive but monostable transcriptional response functions. Using cell sorting and single-molecule footprinting, we found that this variability was heritable and encoded in the local chromatin state of the reporter gene, leading to both reduced synTF binding and lower activation strength when bound. Based on these observations, we developed a quantitative model of gene regulation connecting local chromatin variability to both the probability of a gene being active and its mean expression level once active. Finally^44^, we found that different types of human activation domains can decouple the fraction of active cells in the population from the mean expression level of active cells.

In sum, our results reveal that TFs regulate two separable steps of transcriptional activation (turning a gene from a silent to an active state, then determining the rate of transcription once active) and that stable epigenetic states determine the level of TF input needed to activate expression. These results therefore advance our basic understanding of transcriptional response functions in a minimal expression system in human cells, particularly how TFs interact with chromatin to modulate quantitative features of single-cell and population level responses, and improve our ability to design specific gene expression states for both basic research and therapeutic applications.

## Results

### Transcriptional activation in the Tet-On system is graded at the population level but largely all-or-none at the single-cell level

We first sought to establish a system to measure synTF inputs and gene expression outputs with single-cell resolution in human cells (Figure 1A). We focused our analysis on the Tet-On system (Figure 1B), in which the reverse Tet Repressor (rTetR) is fused to a transactivation domain and binds to Tet Operator (TetO) sequences in the presence of the small molecule doxycycline (dox)^41,43^. We focused on this expression system because it is widely used in many areas of synthetic and cell biology, it is orthogonal to the human genome, and it is inducible and reversible by adding and removing dox^42^. In addition, bimodality has previously been observed in the Tet-On system, similar to other mammalian expression systems^7,14,17,19–21,26^.

To characterize gene expression responses in the Tet-On system, we engineered HEK-293T cells to contain a genomically integrated reporter gene with 9 TetO sites upstream of the commonly used minimal CMV (minCMV) promoter controlling the expression of Citrine (Figure 1B). We inserted this reporter gene at the AAVS1 safe harbor locus using TALENs^45^ to eliminate confounding effects of reporter integration into variable chromatin contexts when measuring input-output functions. To select for correct integrations, we included a splice acceptor and puromycin resistance gene upstream of the TetO sites that is expressed by trapping the upstream PPP1R12C promoter upon successful integration. Analysis of this integration strategy using clonal cell lines revealed a high fraction (>95%) of on-target integrations (Figure S1A). We then engineered a synthetic transcription factor (synTF) by fusing the rTetR DNA binding domain to the fluorescent protein mScarletI3 and VP64, a strong activation domain (AD) containing 4 repeats of the activating region of HSV protein VP16^46^, and placed this construct under the control of the constitutive pEF promoter.

To measure the input-output function of this synTF, we transfected the reporter HEK-293T cells with this construct then added a high concentration (1000 ng/mL) of dox 24 hours after transfection to induce binding at the reporter gene. We then measured the resulting gene expression in single cells 24 hours after dox addition using flow cytometry (Figure 1C). Because the transfection process is stochastic, we were able to generate a wide range of synTF expression levels in the cell population, measured by mScarletI3 fluorescence, and recreate the entire transcriptional response function with a single transfection. Flow cytometry measurements were compensated for fluorescence bleed-through using control transfections with 0 ng/mL dox (Figure S1B) and fluorescence measurements were normalized by forward scatter (FSC) to account for variations in cell size.

From the single-cell fluorescence data, we first calculated the population-level input-output function of the synTF by grouping cells into bins of similar mScarletI3 fluorescence and calculating the average Citrine expression level across cells in each bin (Figure 1C-D). We observed that the population-level input-output function increased continuously with synTF input levels and was well described by a Hill function with a Hill exponent of ∼3.2 (Figure 1D; Figure S1C). To validate this method of measuring transcriptional response functions, we also varied the total amount of synTF transfected into the cells. While increasing the amount of transfected synTF increased the average synTF expression level per cell (Figure S1D), the measured input-output function was robust to the total plasmid amount transfected (Figure 1D; Figure S1E, top). In addition, we perturbed the synTF by replacing VP64 with VP48, which contains only 3 VP16 repeats. We observed that the average transcriptional response function was still continuous and well described by a Hill function but with a lower amplitude, a right-shifted *EC*_50_, and a lower Hill exponent (∼2.4) as expected for a weaker transcriptional activator (Figure 1D, triangles; Figure S1E, bottom).

Despite the continuous transcriptional response function of each synTF at the population level, plotting the entire distribution of Citrine expression levels for each mScarletI3 expression bin revealed that the transcriptional response was largely all-or-none at the single-cell level for rTetR-VP64 (Figure 1E). Within an mScarletI3 bin, cells clearly separated into distinct populations expressing or not expressing the Citrine reporter, producing bimodal gene expression distributions despite all cells having a similar level of synTF expression and a high fraction of cells containing the reporter gene in the same genomic locus. This was also true for the weaker VP48 activation domain, but with less separation between the on and off states (Figure 1E).

Using the single-cell resolution of the flow cytometry data, we decomposed the average transcriptional response functions into the fraction of active cells (Figure 1F) and the mean fluorescence intensity (MFI) of these active cells (Figure 1G). As the synTF abundance increased, the primary response of the cell population was to continuously increase the fraction of cells in the active state (Figure 1F). The average expression level of these cells was also affected by synTF expression level, though to a lesser extent (Figure 1G; compare dynamic range in measured fluorescence to Figure 1D). The responses of both variables to synTF input were right-shifted for VP48 compared to VP64, as shown in Figure 1F and G.

Together these results demonstrate that the continuous population-level responses in the Tet-On synthetic expression system arise predominantly from fractional all-or-none responses at the single-cell level. The primary effect of increasing synTF expression levels is to increase the fraction of cells that activate expression. Notably, this bimodality is present despite the reporter gene being integrated into a genomic safe harbor locus and when only considering cells with similar synTF expression levels. Also of interest, the fraction of cells expressing Citrine appears to be a continuous function of synTF input for both the weaker and stronger activation domains.

### SynTF input regulates both the fraction of active cells and the mean expression level of active cells

We wondered if bimodal gene expression in the Tet-On system was stable over long-term induction of synTF binding or a byproduct of the limited time window available to measure expression following transient transfections of the synTF expression construct (∼48 hours after initial transfection). To investigate the stability of bimodal gene expression distributions, we modified our synTF expression construct and stably integrated it using lentivirus into human K-562 cell lines containing similar reporter genes to those described above (Figure 2A). K-562 cells are well suited to studies of long-term gene expression responses because they are non-adherent and are easily grown in large quantities, allowing us to sample gene expression distributions continuously without significant perturbation (such as repeated rounds of trypsinization).

**Figure 2:**
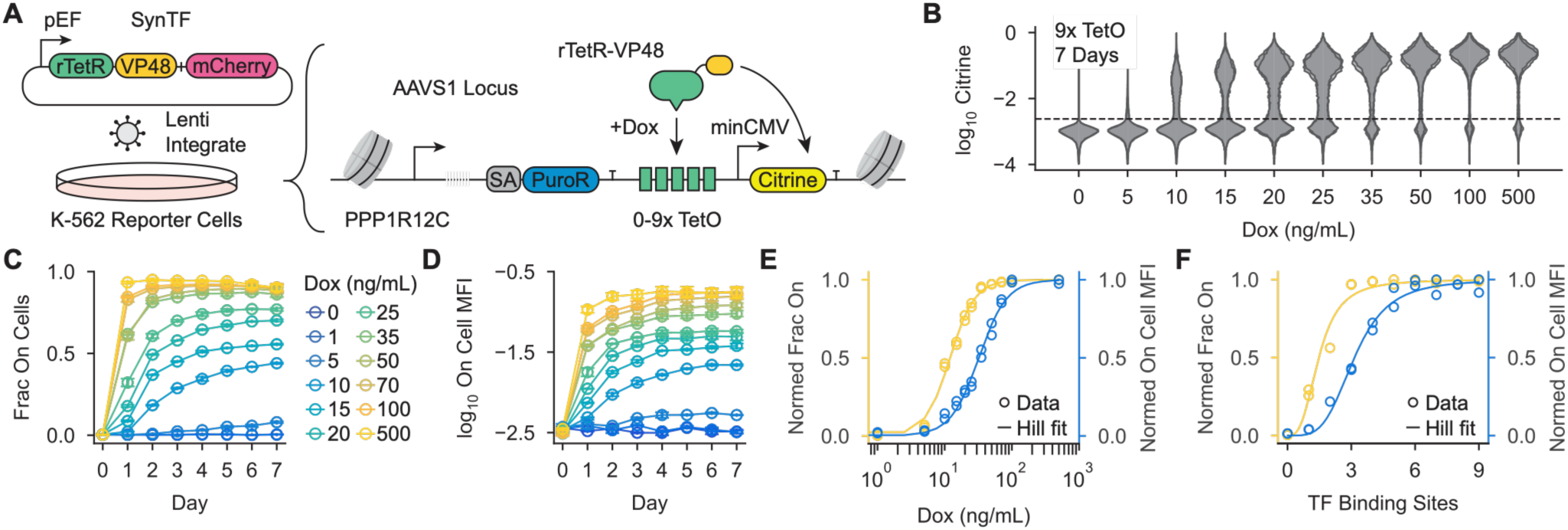
SynTF input controls both the fraction of active cells and their mean expression level over long timescales. A) Synthetic expression system used to quantify transcriptional response functions over long timescales. The synTF consists of rTetR fused to VP48 with mCherry co-expressed from a T2A site and was integrated using lentivirus in K-562 cells. The Citrine reporter gene was integrated into the AAVS1 locus of the cells and selected by promoter-trapping the upstream PPP1R12C promoter upon successful integration. B) Citrine expression distributions following 7 days of synTF recruitment in cells containing a reporter gene with 9 TetO sites at a range of dox concentrations. Two biological replicates are shown. The dashed line represents the gate used to call active cells in C-E. Cells were gated on similar synTF expression levels (Figure S2A-B). C) Fraction of active cells over time during synTF recruitment in cells containing a reporter gene with 9 TetO sites at a range of dox concentrations, as in B. Error is the SD of two biological replicates. D) Mean expression level of active cells over time during synTF recruitment in cells containing a reporter gene with 9 TetO sites at a range of dox concentrations, as in B-C. Error is the SD of two biological replicates. E) Normalized fraction of active cells (yellow) and average expression level of active cells (blue) following 6 days of synTF recruitment in cells containing a reporter gene with 9 TetO sites at a range of dox concentrations (C-D). Values were normalized to the maximum of each variable within biological replicates. Lines are Hill function fits. F) Normalized fraction of active cells (yellow) and average expression level of active cells (blue) following 6 days of synTF recruitment in cells containing reporter genes with 0 to 9 TetO sites using 1000 ng/mL dox. Values were normalized to the maximum of each variable within biological replicates. Lines are Hill function fits.

In these stable cell lines, the synTF is constitutively expressed at a high level. To perturb synTF input at the reporter gene, we either varied the dox concentration used to recruit the synTF (which controls the abundance of synTF available to bind at the TetO sequences) or directly modified the reporter gene by removing TetO sequences from the regulatory region^16^. An mCherry expression marker was co-expressed with the synTF from a T2A site, allowing us to gate over a narrow range of synTF expression levels for downstream analysis (Figure S2A). Over this range of expression levels, we observed minimal correlation between gene expression and mCherry fluorescence, indicating that this method was suitable for quantifying bimodality and mean expression levels (Figure S2B, S2D).

We first recruited rTetR-VP48 in a stable cell line containing 9 TetO sites using a range of dox concentrations for 7 days (Figure 2B). Consistent with our transient transfections, we observed that the cell population split into a bimodal distribution of cells expressing the Citrine reporter gene and cells not expressing the reporter gene and that the fraction of active cells increased with the level of synTF input. At these longer timescales, we also observed a functional relationship between synTF input level and the mean expression level of the active cell population. This indicates that the level of synTF input controls both the probability of a cell being active and the mean transcription rate of each cell when active.

With this stable cell line, we could also carefully measure the dynamics of bimodal gene expression in the cell population. We observed that both the fraction of active cells and their mean expression levels reached steady state by 7 days of synTF recruitment for each dox concentration (Figure 2C-D), indicating that bimodality is stable at the cell population level during long-term synTF recruitment and not caused by cells activating with heterogeneous kinetics. The dynamics of these gene expression distributions are also relatively slow, appearing in the population over a timescale of days at low synTF input (Figure S2E). Because bimodal expression distributions are stable in the cell population, quantitative models of gene regulation that only predict changes in the mean transcription rates, such as commonly used thermodynamic models^28^, do not fully describe transcriptional responses in our model synthetic expression system.

Because synTF input regulates both the fraction of active cells and their mean expression level, we wondered if these two variables had similar dependencies on synTF levels. To compare these two aspects of the transcriptional response on the same scale, we normalized our measurements at each dox concentration to their maximum within biological replicates. We observed that the relationship between dox and each aspect of the transcriptional response was well described by a Hill function, but that they had different dependencies on synTF input (Figure 2E; Figure S2F). The fraction of active cells and their mean expression levels had similar Hill exponents of ∼2.3 and ∼2.1 respectively, but had substantially different *EC*_50_ values (∼12 ng/mL dox and ∼32 ng/mL dox respectively).

To see if these differing dependencies of the fraction of active cells and mean expression level of active cells on synTF input held across different methods of perturbing synTF input, we re-analyzed flow cytometry time courses that we previously collected in stable K-562 lines engineered to have reporter genes with 0 to 9 TetO sites with rTetR-VP48 recruited with 1000 ng/mL dox for several days^16^. We again observed that at low numbers of binding sites the cell population formed stable bimodal gene expression distributions with high and low expressing cells and that the fraction of active cells and mean expression level of active cells increased with the number of binding sites (Figure S2G-I). We fit the relationships between these variables and the number of TetO sites in the reporter gene and again observed that there were substantial differences in *EC*_50_ values (∼1.5 TetO sites and ∼3.2 TetO sites respectively) (Figure 2F; Figure S2J).

Together, these results demonstrate that synTF input levels in the Tet-On system regulate both the fraction of active cells in the population and the mean expression level of these cells, generating long-lasting bimodal gene expression distributions. Interestingly, these variables have separate dependencies on synTF input, with the fraction of active cells saturating at a lower input level than their mean expression.

### Gene expression states are stable over weeks at the single-cell level

It is well established that transcription is stochastic and occurs in bursts of mRNA production interspersed by periods with no transcription^8–10,33,47–54^. To describe this stochasticity, many groups have developed or employed various models of transcriptional regulation where genes are assumed to transition between a number of discrete states, only producing mRNA in certain states or during certain transitions. These include “telegraph” models^8,9,12,22–24,33,49,50,55–60^, kinetic barrier models^8,9,36,61–63^, or various types of transcriptional cycling models^29–32,34,56,64–68^. In principle, the stable bimodal gene expression distributions and all-or-none responses observed for the Tet-On system could be due to bursty transcription of the reporter gene if the dynamics of protein and mRNA degradation are fast compared to the timescales of switching between the gene expression states^22–24^.

To explore this possibility, we analyzed a simple two-state bursting model of transcriptional regulation in which the reporter gene exists in either an active state, where mRNA is transcribed, or a silent state that is impermissive to transcription (Figure 3A; Figure S3A; Methods). Because the Citrine protein used in our reporter gene is relatively stable (a half-life of >24 hours, Figure S3B), switching between gene expression states would have to be on the order of many hours to days to result in bimodal protein expression distributions, which would be much slower than reported switching rates between active and inactive promoter states in mammalian cells^33,47,50^. Therefore, we conceptualized these gene expression states as active and silent “chromatin states,” similar to previously proposed models^12,22^, where transition rates are slower than are typically observed for promoter states. For simplicity, we considered a constant rate of transcription in the active state.

**Figure 3:**
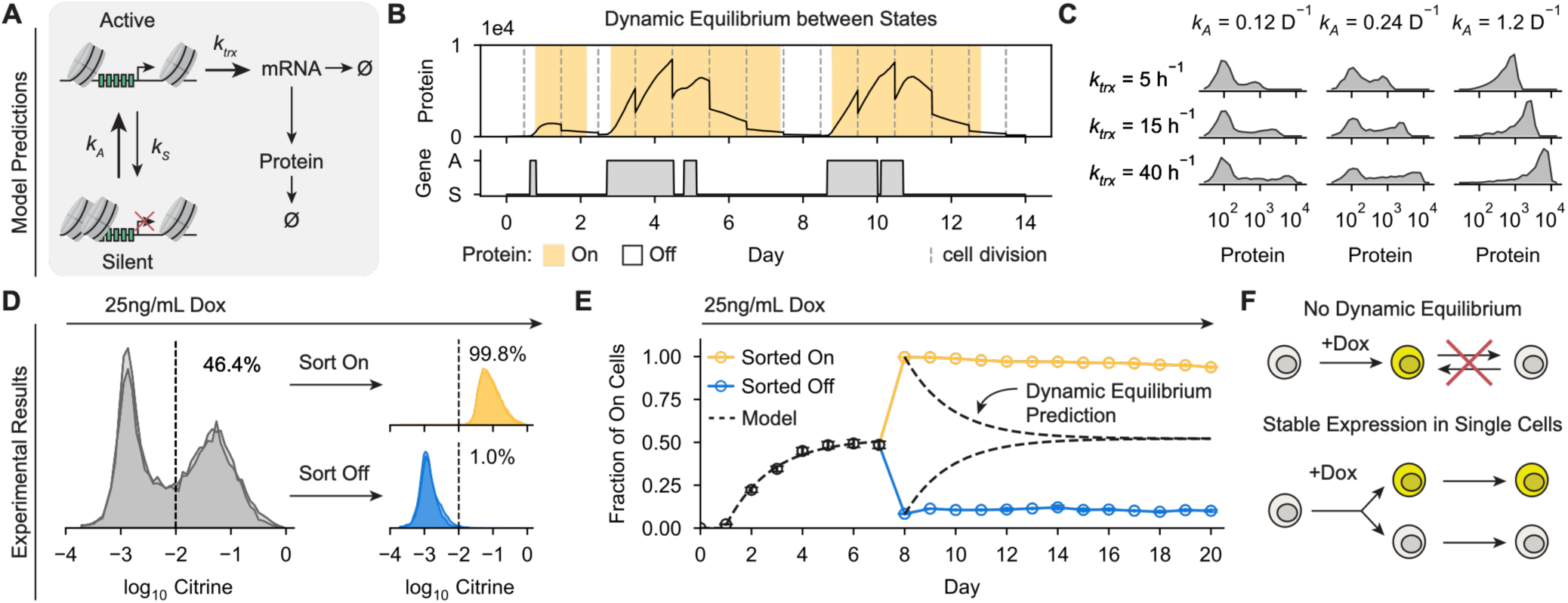
Gene expression states are stable over weeks at the single-cell level and are not the result of a dynamic equilibrium between active and silent gene expression states. A) Two-state bursting model where cells transition between active and silent chromatin states, with transcription only occurring in the active state. B) Example predicted single-cell protein (top) and chromatin state (bottom) dynamics obtained by simulating the model in A using the Gillespie algorithm^77^. A = active state, S = silent state. Cell divisions are indicated by dashed lines. Yellow-shaded regions indicate when a cell would be called as active based on a protein expression threshold, as done in our flow cytometry experiments (here protein > 500). A basal transcription rate of 0.5ℎ^−1^ was assumed in the Silent state to more easily visualize low-expression cells (otherwise, all Off cells would have 0 proteins). Full simulation parameters are shown in Figure S3A. C) Predicted protein expression distributions across populations of 1500 cells obtained by simulating the model in A using the Gillespie algorithm over a range of *k_A_* and *k_trx_* values. Simulations were performed for 14 days of simulated dynamics. Full simulation parameters are shown in Figure S3A. D) Citrine expression distributions in a K-562 cell line expressing rTetR-VP48 and containing a reporter gene with 5 TetO sites before and after cell sorting. rTetR-VP48 was recruited with 25 ng/mL of dox for 7 days (grey), then cells were sorted based on Citrine expression level into On (yellow) and Off (blue) populations and were continued being grown in 25 ng/mL dox. E) Fraction of cells expressing Citrine over time for the pre-sort (black), sorted On (yellow), and sorted Off (blue) cell populations in D. Dashed lines represent the predicted fraction of On cells for each population following cell sorting assuming a dynamic equilibrium between an active and silent chromatin state and fitted on the pre-sort fraction of On cells for days 0-7 (Figure S3D-E). Error bars represent the SD of two biological replicates. F) Gene expression states are stable at the single-cell level, ruling out a dynamic equilibrium between active and silent gene expression states as the underlying cause for bimodality.

Under this model, individual cells stochastically transition between the active and silent chromatin states on a slow enough timescale that the protein produced in the active state is likely to be fully degraded during the subsequent silent periods (Figure 3B). Sampling a population of cells at a single timepoint would therefore result in a bimodal distribution of gene expression roughly corresponding to the fraction of active and silent cells at that time point (Figure 3C). Intriguingly, we found that in this model the fraction of active cells can be continuously tuned by the rate *k_A_* in a manner largely independent of the mean expression level of active cells, which is controlled by mRNA and protein production/degradation rates (Figure 3C). A TF functioning simultaneously on these processes would therefore tune both the fraction of active cells and their mean expression level but not necessarily with the same sensitivity, as observed in our experimental results (Figure 2).

Fundamentally, bimodality in this class of gene regulatory model is the result of a dynamic equilibrium between active and silent cells with protein production and degradation rates fast enough to efficiently clear proteins in the amount of time that cells spend in the silent states. To test if a bursting phenomenon was responsible for bimodality in the Tet-On system, we generated a steady state bimodal gene expression distribution in K-562 cells by recruiting rTetR-VP48 with a low dox concentration (25 ng/mL), and then sorted cells by gene expression state. We then continued culturing these sorted populations in the same low dox dose (Figure 3D; Figure S3C). A bursting model would predict that the sorted cell populations would re-equilibrate to the pre-sort steady state fraction of active cells over a similar timescale as the original approach to steady state (Figure 3E, dashed lines; Figure S3D). However, sorted On and Off cell populations did not re-equilibrate during 12 days of synTF recruitment after cell sorting (Figure 3E).

Together, these results demonstrate that bursting is not responsible for the bimodality observed in the Tet-On system despite bursting models being able to describe well many quantitative features of transcriptional responses in this context. Instead, gene expression states in this system are long-lived at the single-cell level and are therefore not the result of a dynamic equilibrium between active and silent gene expression states (Figure 3F), as is assumed by these models.

### Variability in ultrasensitive, but monostable, transcriptional responses produces bimodal expression distributions

Given these findings, we wondered what underlying mechanism is responsible for the continuous responses at the population level, all-or-none responses at the single-cell level, and stability of single-cell gene expression states. At intermediate synTF levels, our results suggest that a subset of cells are stably induced to a high expression state while the remaining cells in the population remain in an inactive state, analogous to *Xenopus* oocytes maturing in an all-or-none manner in response to increasing progesterone concentrations^69^. We reasoned that two potential underlying mechanisms are consistent with our observations: (1) bistability at the single-cell level mediated by a feedback mechanism with cell-to-cell variability in the amount of synTF input needed to engage feedback loops, similar to *Xenopus* oocyte maturation, or (2) ultrasensitive but monostable transcriptional responses that vary between individual cells and where the variations persist over a timescale of weeks (Figure 4A).

**Figure 4:**
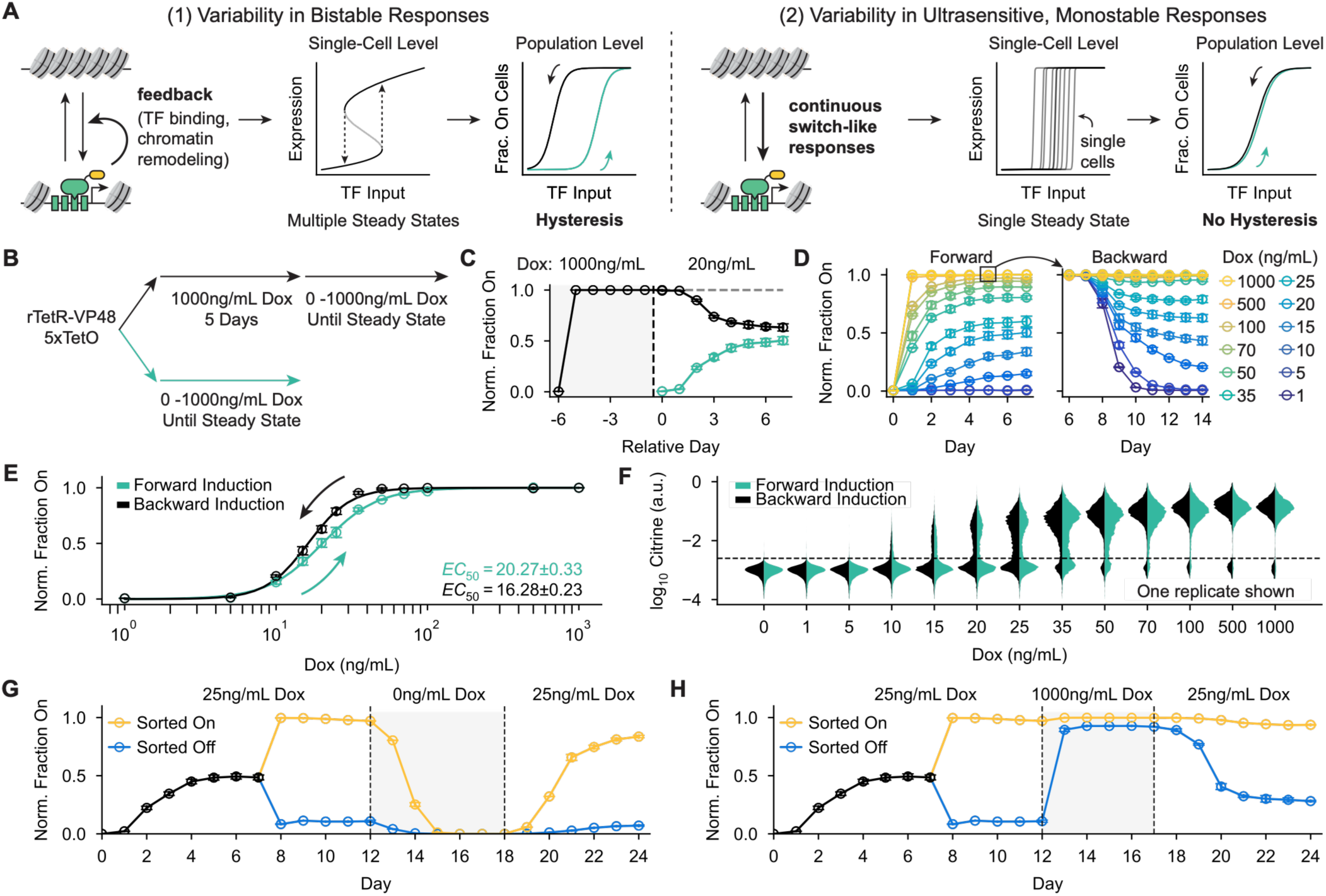
Variability in ultrasensitive, but monostable, transcriptional responses produces bimodal expression distributions. A) Potential models that could explain the observed all-or-none gene expression responses and the single-cell stability of responses in the Tet-On system: (1) bistability from transcriptional feedback, which would result in hysteresis as the amount of synTF is reduced, or (2) ultrasensitive but monostable responses where individual cells require variable amounts of synTF to activate gene expression, which would not produce hysteresis. B) Experimental design to measure hysteresis in the Tet-On system. For the forward induction, reporter cells expressing rTetR-VP48 were cultured in different dox concentrations until steady state (teal line). For the reverse induction, the same cell line was first cultured in 1000ng/mL dox for 5 days, washed in PBS, and then re-cultured in lower dox doses until steady state (black line). C) Normalized fraction of reporter cells expressing Citrine over time during forward (teal) and backward (black) inductions of the synTF to a final concentration of 20 ng/mL dox. Normalization procedure is described in Figure S4C-E. Error bars represent the SD of two biological replicates. Vertical dashed line represents the time of dox downshift. D) Normalized fraction of reporter cells expressing Citrine over time during forward and backward inductions of the synTF with varying dox concentrations, as described in B. Normalization procedure is described in Figure S4C-E. Error bars represent the SD of two biological replicates. E) Steady state fraction of cells expressing Citrine at various dox concentrations during forward and backward inductions (from D). Hill function fits (solid lines) were used to estimate the *EC*_50_ of each induction (uncertainty is the standard deviation of each fit). Error bars represent the SD of two biological replicates. F) Citrine expression distributions at various dox concentrations during forward and backward inductions from C-E. G) Normalized fraction of active cells over time during dox washout and re-addition for sorted populations (same initial sort as in Figure 3B-E). After 3 days of induction at 25 ng/mL dox, sorted cells were washed with PBS and re-suspended in 0 ng/mL dox (grey shading). After 6 days, 25 ng/mL dox was re-added. Error bars represent the SD of two biological replicates and measurements were normalized relative to the no-washout control. H) Normalized fraction of active cells over time during transient dox upshift for sorted cell populations (same initial sort as in Figure 3B-E). After 3 days post-sort, cells were washed with PBS and re-suspended in 1000 ng/mL dox media (grey shading). After 5 days, sorted cells were washed with PBS and re-suspended in 25 ng/mL dox media. Error bars represent the SD of two biological replicates and measurements were normalized relative to the no-upshift control sort.

Gene expression states and DNA accessibility are associated with certain chromatin modifications^12,70–73^ and are often regulated through reader/writer enzymes that can be mutually antagonistic and have the capacity for feedback^74^. Positive feedback of active transcription (e.g. a TF binding then recruiting chromatin-opening enzymes which allow more TFs to bind) could in principle lead to bistability in gene expression states (as previously proposed^26^) resulting in stable, all-or-none transcriptional responses at the single-cell level^75^. If there were cell-to-cell variability in the amount of TF necessary to evict nucleosomes and engage this positive feedback loop, the transcriptional response would appear graded and bimodal at the population level, with the degree of variability determining how graded the population-level response is (Figure 4A, left).

Alternatively, the ultrasensitive monostability model posits that only a single stable gene expression state exists at each synTF input level. If transcriptional responses to synTF inputs are sufficiently ultrasensitive, small changes in synTF inputs would create effectively all-or-none responses in single cells. In addition, if the midpoint of the transcriptional response function varied between cells, only a subset of cells would be activated at intermediate synTF concentrations. This mechanism would therefore predict bimodal, graded responses at the population level but stable, effectively all-or-none responses at the single-cell level (Figure 4A, right). The sharpness of the population-level response would be determined by the degree of variability in single-cell response functions.

A key difference between these models is that bistability would predict hysteresis, or even irreversibility, in the fraction of active cells: once in the active state, positive feedback allows for a cell to more easily remain in the active state even if synTF input is lowered. Notably, the high levels of stability we observed between the sorted cell populations (Figure 3E) suggest that the separation between the stable gene expression states is much larger than the noise or fluctuations in the system (i.e. stochastic transitions between these steady states are very rare events). The alternative model, where transcriptional responses vary between cells and are ultrasensitive but monostable, would not predict hysteresis.

To distinguish between these possibilities, we took advantage of the reversibility of rTetR binding to systematically quantify hysteresis in this system. We induced K-562 cells expressing rTetR-VP48 and containing a similar genomically integrated reporter gene to that described above using dox concentrations ranging from 0 to 1000 ng/mL. We refer to this initial induction as the forward induction. After 5 days of induction, we took a subset of the maximally induced cells (1000 ng/mL dox), washed them, and replated them in the same range of dox concentrations as the initial induction (Figure 4B). We refer to this downshift in dox concentration as the backward induction. This workflow allowed us to compare the steady state fraction of active cells in the population at the same dox concentration between the forward and backward inductions. Looking at a single dox concentration (20 ng/mL), we observed that the forward and backward inductions resulted in very similar steady state fractions of active cells (Figure 4C).

We then carefully measured the steady state fraction of active cells during forward and backward inductions for a range of dox concentrations and normalized the data to account for background silencing of the reporter gene over the induction time course (Figure 4D; raw data in Figure S4A-B; normalization procedure in Figure S4C-E). When comparing the steady state fraction of active cells for each level of synTF input, we observed only a small left-shift of the population-level response function for the backward induction relative to the forward (∼16 ng/mL dox vs ∼20 ng/mL dox respectively; Figure 4E; Figure S4F). The presence of a small amount of hysteresis is not surprising given the prevalence of feedback in chromatin regulation. Alternatively, our forward or backward inductions might not quite be in steady state at the time of measurement. In either case, the measured level of hysteresis is too small to account for the observed stability of single-cell gene expression states, or for the bimodality at the population level. Indeed, when we compared full Citrine expression distributions for each dox dose between the forward and backward inductions, we observed that at intermediate synTF inputs the cell population appeared highly bimodal even for the backward induction, with a large fraction of cells turning off Citrine expression entirely (Figure 4F). These results support a model where long-lived cell-to-cell variability in ultrasensitive, monostable transcriptional responses results in bimodal gene expression distributions in the Tet-On system rather than bistability from potential chromatin-mediated positive feedback.

To further test whether cell-to-cell variability in transcriptional response functions was responsible for the observed bimodal gene expression distributions, we returned to the sorted On and Off cell populations cultured in 25 ng/mL dox (Figure 3E). We took a split of each sorted cell population and washed off the dox for 6 days. We then re-induced the cells with 25 ng/mL of dox and observed that the sorted Off cells largely failed to activate while the majority of the sorted On cells re-activated (Figure 4G; Figure S4G). These results indicate that the cells do not return to a shared “ground state” in the absence of dox and that the variability between cells persists for many days and cell divisions.

Conversely, we took the same sorted cell populations at 25 ng/mL dox, and instead of washing out dox, increased the dox concentration to 1000 ng/mL for 5 days before washing the cells and replating in 25 ng/mL dox (Figure 4H; Figure S4H). During the transient treatment with 1000 ng/mL dox, both cell populations almost entirely activated gene expression. Following the return to 25 ng/mL dox, almost all of the sorted On cells remained in the active state, while the sorted Off cells largely turned off Citrine production, with a slight increase in the fraction of active cells that was consistent with our previous hysteresis measurement. These results reveal that individual cells in the population have different synTF thresholds for activating gene expression in the Tet-On system, which gives rise to bimodal expression distributions, but these ultrasensitive transcriptional response functions are not caused by bistability. Moreover, these thresholds are unchanged by transcriptional activation.

Finally, we sorted individual clones from our K-562 reporter line and activated these clonal lines with low (25 ng/mL) and high (1000 ng/mL) dox. We observed that, while all clonal lines activated at high dox, only ∼50% of the clones activated at low dox and this activation was all or none in nature (Figure S4I-J), consistent with our model that heritable cellular variability in ultrasensitive transcriptional response functions results in bimodal gene expression distributions in the cell population. These data also highlight that variability in transcriptional responses is heritable and stable over many cell generations, as individual clones were expanded for ∼2 weeks before recruitment assays were performed.

Together, these experiments do not support bistability as the underlying cause of the all-or-none single-cell gene expression responses in the Tet-On system. Instead, they support a model where individual cells have highly ultrasensitive, but monostable, input-output functions that have variable switching points across the population, leading to bimodal gene expression distributions. Given the observed continuous relationship between synTF input and the fraction of active cells, it seems likely that this variability must create a relatively continuous distribution of “activation potentials” for cells in the population. In this case, modest increases in synTF input would result in a comparable increase in the number of cells that pass their specific synTF activation threshold and transition to the active state, resulting in a graded population level response.

### Variability in transcriptional activation is largely encoded at the locus rather than at the cell level

After finding that bimodality in the Tet-On system is the result of heritable variability between ultrasensitive, monostable transcriptional response functions, we asked what the source of this variability was in the cell population. In principle, it could be encoded at the cellular level in *trans* (e.g. the expression levels of coactivators, signaling states, etc.), in *cis* (e.g. the local chromatin environment or histone modifications of the reporter gene), or a combination of the two.

To quantify the relative contributions of cell-level and locus-level effects on the heterogeneity of transcriptional responses, we carried out experiments on cells expressing two reporter genes in separate copies of the same genomic locus. We designed reporter genes with 4 TetO sites upstream of the minCMV promoter driving expression of either Citrine or mCherry (Figure 5A). We integrated both reporters into K-562 cells targeting the AAVS1 locus, then stably integrated a rTetR-HALO-VP48 construct in these dual-reporter cells using lentivirus. Finally, after puromycin selection, we transiently added dox to the cells and sorted a pure population that expressed both reporter genes. The sorted cells expressing both genes would thus contain the reporters integrated into copies of the AAVS1 locus on separate chromosomes. For expression measurements, we stained cells with HALO dye (JF-646) to gate cells with similar synTF expression levels for downstream analysis (Figure S5A). We induced gene expression in this cell line at a range of dox concentrations for 6 days and verified that both reporters activated to similar levels in the cell population (Figure S5B).

**Figure 5:**
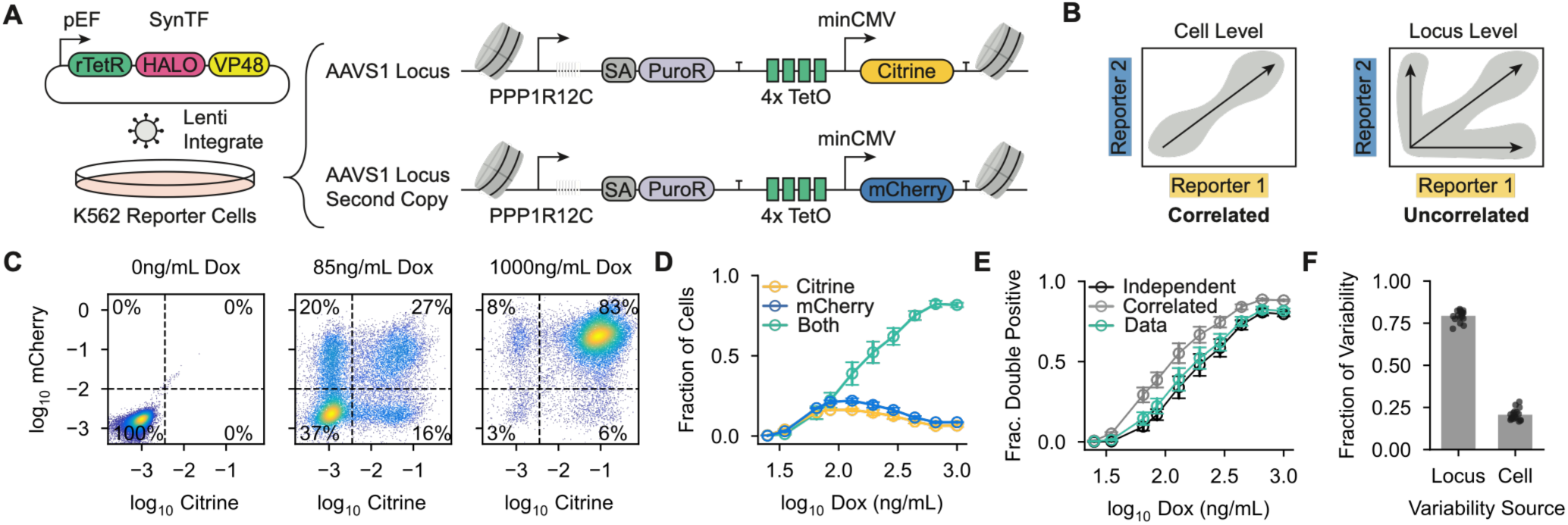
Variability in transcriptional activation is largely encoded at the locus rather than at the cell level. A) Engineered dual-reporter system for measuring correlated gene activation in the Tet-On system. Citrine and mCherry reporter genes were co-transfected into K-562 cells and integrated into copies of the AAVS1 locus. The synTF was fused to HALO-tag and lentivirally integrated in the reporter cells. After puromycin selection, cells were transiently induced with 100 ng/mL dox and cells expressing both reporters were purified using cell sorting. B) Expected gene expression between reporters if the variability in transcriptional response functions was encoded at the cell level (correlated expression) or locus level (uncorrelated expression). C) Single-cell Citrine and mCherry expression after 6 days of synTF recruitment at different dox concentrations in the dual-reporter cell line described in A. Points are colored by density. Cells were gated on synTF expression level for downstream analysis. Dashed lines represent gates used to call gene expression states. Percentages are the proportion of cells in each quadrant. D) Fraction of cells expressing Citrine only, mCherry only, or both Citrine and mCherry after 6 days of synTF recruitment at a range of dox concentrations (quantified from C). Error bars represent the SD of two biological replicates. E) Comparison between fraction of cells expressing both Citrine and mCherry (teal) and the expected fraction of double-positive cells if each reporter were entirely independent of the other (black) or perfectly correlated (grey). Error bars represent the SD of two biological replicates. F) Decomposition of variability in transcriptional activation into locus-level and cell-level contributions (Methods) from the measurements in E (Figure S5E-F). Error bars represent the SD of two biological replicates. Points represent calculations from individual dox doses.

If cell-level *trans* effects were responsible for the observed heterogeneity, we would expect highly correlated expression between the two reporters (Figure 5B, left). Conversely, if locus-level *cis* effects were responsible, we would expect gene activation to be largely uncorrelated between the reporters (Figure 5B, right). From single-cell Citrine and mCherry measurements, it was apparent that many cells only expressed one of the two reporters at low dox doses after 6 days of synTF recruitment, and at maximal dox induction most cells in the population expressed both reporters (Figure 5C; Figure S5C). We quantified the fraction of cells in each gene expression state (Citrine only, mCherry only, both reporters, or neither) for each dox concentration and observed that the fraction of cells in the mCherry-only and Citrine-only populations peaked at low dox levels before decreasing as more cells moved to the double-positive state (Figure 5C, D). This is consistent with a model where locus-level variability in the amount of synTF needed for transcriptional activation results in discordant expression between the reporters at low input levels. At higher synTF input levels, gene expression becomes more uniform in the population as the activation threshold for both reporters is reached in more cells.

From the relative abundance of each gene expression state in the cell population, we then sought to quantify the independence of the reporters in individual cells. As a first pass, we compared the measured fraction of double-positive cells to the predicted fraction of double-positive cells assuming complete independence (multiplying the fractions of Citrine and mCherry positive cells across the entire population) vs. complete concordance (every cell expressing one reporter expresses the other as well) between the reporters (Figure 5E). The observed fraction of double-positive cells was notably more similar to the prediction from the independence model across all tested dox doses, but exhibited a small but consistent enrichment of double-positive cells over the independent expectation (Figure 5E; Figure S5D). We then quantified the relative contribution of locus-level and cell-level variability in activation across dox doses assuming identical activation of the two reporters, as done previously to analyze intrinsic vs. extrinsic noise in gene expression (Figure S5E-F)^76^. Using this method, we determined that approximately 75% of the variability in transcriptional response functions is encoded in the local environment of the reporter gene, while 25% originates from more global cell-level effects (Figure 5F). Notably, this variability must be inherited through many rounds of cell division, persist throughout active gene expression states, and encode a nearly continuous distribution of “activation potentials” across the cell population. These features suggest that variability in transcriptional responsiveness is primarily encoded in the local chromatin environment of the transgene through an epigenetic mechanism.

### Local variability is associated with reductions in both synTF binding and the ability of the synTF to activate gene expression when bound

Given that the variability in single-cell transcriptional response functions is primarily caused by heterogeneity in the local environment of the reporter gene, we wondered what aspects of synTF function differed in cells with high vs. low activation potentials. To answer this question, we used a recently developed version of amplicon-based single-molecule footprinting (SMF) to measure local accessibility and synTF binding at the reporter^35^. In this method, isolated nuclei are treated with the recombinant methyltransferase M.CviPI, which then methylates accessible cytosines in a GpC context. Unmethylated cytosines are then enzymatically converted to uracils and single-molecule accessibility states can be read out with next-generation sequencing (Figure 6A). Comparing the sizes of methylation footprints and their locations along an annotated amplicon allows for the quantification of synTF binding levels and nucleosome occupancy in the cell population^35^.

**Figure 6:**
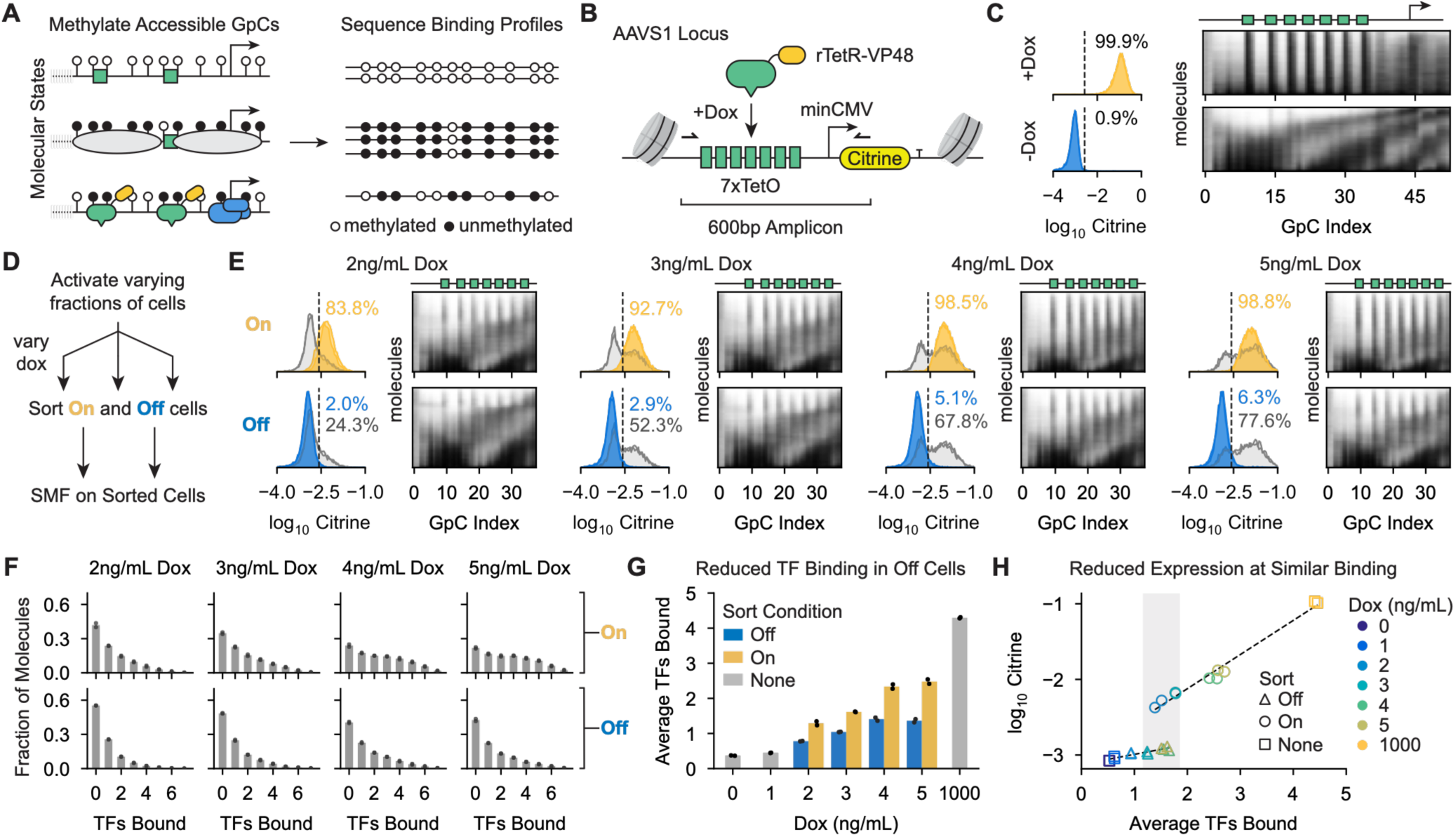
Local variability is associated with reductions in both synTF binding and the ability of the synTF to activate gene expression when bound. A) Schematic of reading molecular states with SMF. Left: isolated nuclei were treated with the M.CviPI methyltransferase, which methylates accessible Cs (white circles) in a GpC context. Right: unmethylated Cs (black circles) are then enzymatically converted to Us before amplification and read out as Ts during sequencing. Grey ovals represent nucleosomes, blue complex represents transcriptional machinery. B) Reporter system for performing amplicon SMF. A reporter gene with 7 TetO sites upstream of the minCMV promoter and containing a high density of GpC sites was integrated into the AAVS1 locus of K-562 cells constitutively expressing rTetR-VP48, as previously described. A 600bp region capturing the TetO sites and minCMV promoter was amplified for sequencing following methylation and enzymatic conversion. C) Example single-molecule footprints for cells treated with 1000 ng/mL dox (top) and 0 ng/mL dox (bottom). Citrine expression distributions are shown on the left, and corresponding single molecule methylation profiles are shown on the right. Each row of the heatmap is an individual molecule, each column is a GpC site in the reporter amplicon. White = methylated/accessible, Black = unmethylated/protected. Positions of TetO sites (green boxes) and the TATA box (arrow) are shown above. D) Experimental design for SMF experiment. Cells containing the SMF reporter were treated with varying concentrations of dox for 11 days to activate different fractions of cells, then sorted into Off and On populations. The following day, SMF was performed on each sorted population. E) Citrine expression distributions before (grey) and after (yellow = On, blue = Off) cell sorting after 11 days of VP48 recruitment at various dox concentrations, with corresponding single molecule footprints for each condition (right). Each row is an individual molecule’s methylation profile; each column is a GpC site in the reporter amplicon. White = methylated/accessible, black = unmethylated/protected. Positions of TetO sites (green boxes) are shown above. F) SynTF binding distributions for the dox concentrations and sort conditions shown in E. Black points indicate biological replicates for each condition. Binding distributions were calculated using a maximum likelihood model, as previously described^35^. G) Average number of synTFs bound for the conditions and binding distributions in F. Dots indicate biological replicates. H) Relationship between average number of TFs bound (from G) and average Citrine expression level for each dox concentration and sort condition. Lines are linear fits to log MFI in each regime to illustrate trends.

We performed SMF on a genomically integrated reporter similar to that described above but modified to contain a high density of cytosines in GpC contexts for improved resolution of molecular binding states (including TetO sequences flanked by GpCs to infer synTF binding). After methylation and enzymatic conversion, we amplified a 600bp region of the reporter gene containing all TetO sites and the minCMV promoter region to analyze the reporter locus (Figure 6B). As a validation for this method, we first compared methylation profiles in K-562 cells treated with 1000 ng/mL dox or 0 ng/mL dox. We sorted molecules by average accessibility for each condition and generated heatmaps where each row corresponded to an individual molecule and was colored according to methylation status (white = methylated, black = unmethylated), shown on the right of Figure 6C. In the 1000ng/mL dox condition, we observed prominent stripes of protection that corresponded to the positions of TetO sequences in the amplicon, indicating strong binding and footprinting of the synTF (Figure 6C, top; Figure S6A). These stripes were largely absent in the 0 ng/mL dox condition. Instead, we saw broader regions of protection that roughly corresponded to the size of nucleosomes and were phased across the reporter (Figure 6C, bottom; Figure S6A).

After validating that we could measure synTF binding with SMF, we sought to use this method to measure differences in synTF occupancy and activity between cells with different levels of activation potential. To do so, we treated reporter cells with varying dox doses for 11 days, producing steady state gene expression distributions with different fractions of active cells (Figure S6B). We then sorted active and silent cells for each dox dose and performed SMF on each sorted population (Figure 6D-E). To ensure that methylation and conversion efficiencies were similar across all conditions, we spiked in a cell line containing a reporter gene with a single CTCF binding site and found that this motif footprinted similarly across conditions (Figure S6C). We also sorted cells on synTF expression level to ensure a pure cell population served as input to the footprinting protocol (Figure S6D).

With this experimental design, we generated a range of On and Off gene expression states, but with the cells in each state requiring a different synTF input level to activate. For example, cells that do not activate when treated with 5 ng/mL dox (where ∼75% of the population activates) have a lower activation potential on average than cells that do not activate when treated with 2 ng/mL dox (where only ∼25% of the population activates) (Figure 6E). In addition, the mean expression level of the sorted On populations increased with dox dose, allowing us to investigate the relationship between synTF binding and the pre-sort fraction of On cells, the MFI of the On population, and the gene expression state (On vs Off) (Figure 6E).

After performing SMF on each sorted cell population at each dox concentration, we noticed clearly visible stripes of synTF binding at the TetO motifs (Figure 6E) compared to the -dox control (Figure 6C) even in steady-state Off populations. To more precisely quantify levels of synTF binding at the reporter gene across the various conditions, we used a previously developed probabilistic model that identifies the most likely configuration of synTF binding and nucleosome positions for a given methylation profile^35^ (see Figure S6E for example molecules and assigned states). This model allowed us to infer the distribution of the number of synTF molecules bound at the TetO sites in each condition (Figure 6F; Figure S6F). We observed that within both the On and Off sorted conditions, the synTF binding distribution shifted right (to higher synTF binding levels) as the dox concentration increased (Figure 6F, rows; Figure 6G). In addition, within a single dox concentration, the synTF binding distribution was left shifted (to lower synTF binding levels) in sorted Off cells compared to sorted On cells (Figure 6F, columns; Figure 6G).

We then calculated the average number of synTFs bound in each cell population and compared this value across dox concentrations and gene expression states (Figure 6G). We again observed that the average synTF binding increased in both On and Off cells as dox increased, but there was systematically lower synTF binding in the Off cells compared to the On cells. In addition, the difference in average synTF occupancy between the On and Off cells increased as a function of dox dose.

We wondered how average synTF binding levels related to the mean expression level of each cell population and if average binding was sufficient to predict whether or not a cell population was On or Off regardless of dox concentration. Surprisingly, when we compared these variables, we observed a regime where cell populations had nearly identical average synTF binding levels but opposite gene expression states (Figure 6H, shaded region). For example, sorted Off cells treated with 4 or 5 ng/mL dox had the same average binding, ∼1.5 synTFs, as sorted On cells at 2 ng/mL dox. Once gene expression was activated, the mean expression level of the cell population increased exponentially with average synTF binding levels (Figure 6H).

Together, these results demonstrate that heterogeneity in the local chromatin environment of the reporter gene introduces variability in transcriptional response functions through multiple mechanisms. For a single synTF induction level, a subset of cells in the population have reduced synTF binding at the reporter gene and fail to activate gene expression. In addition, cells that are harder to activate in the population need more synTF binding to activate gene expression than cells that are easier to activate, indicating that local heterogeneity reduces the activation ability of bound synTFs in a subset of cells.

### Gene activation in the Tet-On system can be quantitatively described by a two-step model with locus-level heterogeneity

Based on our experimental results, we wondered if we could incorporate chromatin-mediated heterogeneity into a quantitative model of transcriptional regulation to better account for the single-cell and population-level responses we observe in the Tet-On system. Since we found that synTF input regulates both the probability of a cell being active and the mean expression level of active cells (Figure 2), we implemented a two-step model where cells can transition from a silent to active gene expression state with a rate constant *k_A_* and transcription occurs in the active state with a rate constant *k_trx_* (Figure 7A). Because we observed that transitions from the silent to active state are reversible (Figure 4) we included a back rate from the active to silent state with the rate constant *k_S_*. Together, these transitions define a monostable system with a unique steady state probability of being active for each value of *k_A_* and *k_S_*.

**Figure 7:**
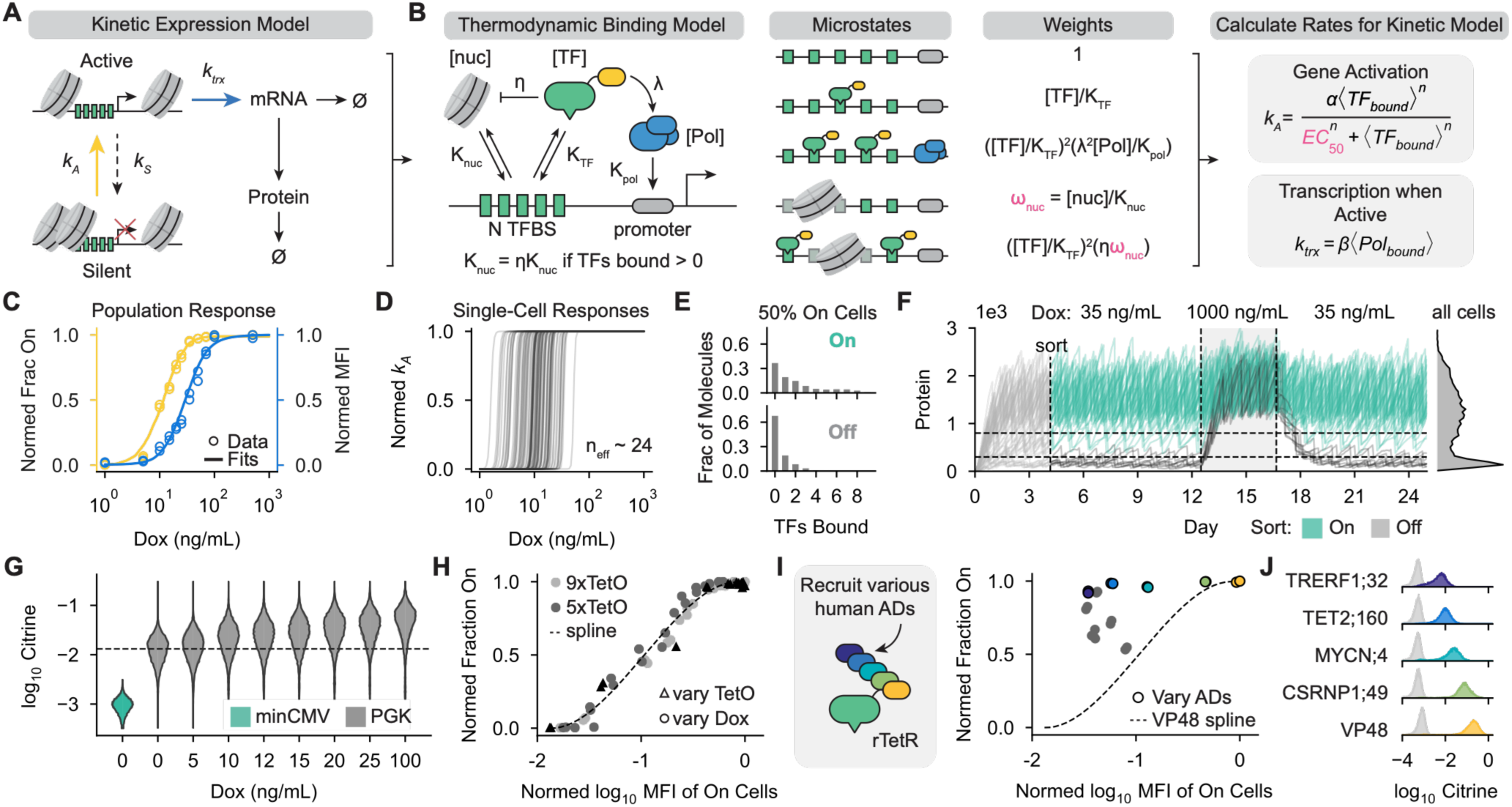
Gene activation in the Tet-On system can be quantitatively described by a two-step model with locus-level heterogeneity, and activation domains can decouple the fraction of active cells and their mean expression level. A) Two-state kinetic model used to simulate single-cell gene expression dynamics. B) Thermodynamic binding model used to calculate the gene activation and transcription rates. SynTFs compete with and remodel nucleosomes to bind to the TetO array, as previously described^35^, and stabilize the binding of RNA polymerase to the promoter. The average occupancy of the synTF and RNA polymerase are calculated from the partition function, and then used to calculate the activation and transcription rates for the kinetic model in A. Parameters with variability between cells are colored pink. C) Model activation and transcription rate fits to normalized fractions of active cells (yellow) and mean expression levels of active cells (blue) at various dox concentrations, from Figure 2E. Cells with a normalized *k_A_* > 0.9 were considered On. *ω_nuc_* and the *EC*_50_ of *k_A_* were sampled for 300 cells from a fit multivariate lognormal distribution with a log-space correlation coefficient of 0.6 between the parameters (Methods). D) 100 sample predicted single-cell transcriptional response functions used to calculate the population level responses in C. The effective Hill exponent was calculated empirically for each cell^75^. E) Predicted synTF binding from the model and fits in A-D. An effective synTF concentration of 14 ng/mL dox was used, then On (*k_A_* > 0.9) and Off cells were separated to calculate predicted synTF binding distributions. F) Single-cell gene expression dynamics^77^ predicted using the model and parameters in A-D. Dox concentrations used are indicated above each region. After 100 hours, cells with >800 proteins were designated as On (teal) and cells with <300 proteins were designated as Off (black). Within the shaded region TF input was transiently increased from 35 to 1000 ng/mL dox. The gene expression distribution on the right indicates the protein expression distribution in the entire population of cells after 25 days. Trajectories for 50 cells are shown. G) Gene expression distributions after 4 days of rTetR-VP48 recruitment in K-562 reporter cells containing an AAVS1-integrated reporter gene with the 9xTetO-PGK promoter (grey), instead of the 9xTetO-minCMV promoter (teal), at various dox doses. Dashed line is the mean PGK expression level with 0 ng/mL dox. H) Relationship between fraction of active cells and their mean expression level across several experiments perturbing rTetR-VP48 input at the minCMV promoter in various ways. A spline fit is shown as a dashed line. I) Relationship between the normalized fraction of active cells and their mean expression level after 5 days of recruitment for synTFs with different activation domains at 1000 ng/mL dox at the 9xTetO-minCMV promoter. The dashed line represents the spline fit for rTetR-VP48 from H. Colored points correspond to domains highlighted in J, grey points correspond to other domains tested. MFI measurements were normalized to the expression level induced by rTetR-VP48 at 1000 ng/mL dox. J) Citrine expression distributions (left) and average Citrine expression levels (right) for 5 synTFs with different activation domains produced after 5 days of recruitment with 1000 ng/mL dox at the 9xTetO-minCMV promoter. Grey histograms indicate AD-matched no dox controls, colored histograms represent cells treated with 1000 ng/mL dox.

Because slow, dynamic transitions between gene expression states could not explain bimodality in this system, we considered the regime where, if synTF input is over a certain threshold, the rate of activation is much greater than the rate of silencing (*k_A_* ≫ *k_S_*) and both rates are fast enough to generate unimodal expression distributions. When synTF input is below this threshold, the rate of activation is much less than the rate of silencing (*k_A_* ≪ *k_S_*). We therefore hypothesized that, for a single-cell, the activation rate *k_A_* is a highly ultrasensitive function of synTF input levels. Once in the active state, *k_trx_* is also regulated by synTF input but separately from the regulation of *k_A_*, as observed experimentally.

We then sought to connect this kinetic model to the effects of local chromatin heterogeneity on the function of synTFs, based on our footprinting data (Figure 6). To do this, we adapted a recently proposed thermodynamic model of rTetR-VP48 binding^35^ where synTFs compete with nucleosomes for an array of binding sites and, when bound, inhibit nucleosome binding through an unfavorable energetic interaction (Figure 7B, left). To model rates of transcription, we incorporated RNA polymerase binding to the promoter in the model and represented the regulatory effect of synTFs through favorable interactions between them and polymerase when co-bound.

This framework allowed us to calculate synTF and polymerase binding by enumerating all molecular states and their associated statistical weights based on binding/interaction energies and the effective concentration of the synTF, our most accessible experimental parameter (Figure 7B, middle). To connect our kinetic expression model with these molecular states, we assumed *k_trx_* was proportional to the average occupancy of RNA polymerase (the “occupancy hypothesis”)^9,28^ and assumed that *k_A_* was related to the average occupancy of the synTF through a Hill function (Figure 7B, right; Figure S7A). At its core, this combined thermodynamic and kinetic framework allowed us to relate synTF concentration and the molecular features of the reporter gene (i.e., nucleosome occupancy and number of binding sites) to the average binding of the synTF, and then connect average binding to both the probability of the reporter gene being active and its mean expression level (Figure S7B). We found it necessary to decouple the dependencies of *k_A_* and *k_trx_* on synTF occupancy because of our observations that the fraction of active cells and mean expression level of active cells saturated at different input levels (Figure 2).

Finally, we incorporated chromatin-encoded heterogeneity into our model by assuming that it introduces variability in nucleosome binding (using the parameter *ω_nuc_*) and the activation strength of the synTF when bound (through the *EC*_50_ of *k_A_*) (Figure 7B, pink parameters). We chose to represent the variability in response functions in this way because (1) most of the variability was encoded in the local environment of the reporter gene (Figure 5); (2) there was a systematic reduction in synTF binding in Off cells compared to On cells (Figure 6G); (3) there was a negative relationship between inferred nucleosome binding at the TetO array and synTF binding (Figure S7C); and (4) there was a reduction in the ability of the synTF to activate gene expression in Off cells even when bound at comparable levels to On cells (Figure 6H). In addition, we assumed that these parameters were stable at the single-cell level due to how long-lived the variability in transcriptional response functions was in our experimental observations (Figure 4) and that each simulated cell contained one reporter gene. For a full model description and parameter values, see Methods.

We then fitted this model to our data (Figure S7D) to see if our framework and assumptions were sufficient to recapitulate the totality of our experimental observations. Briefly, we simulated heterogeneity in individual cells by sampling nucleosome binding parameters and the *EC*_50_ of *k_A_* from a multivariate log-normal distribution, then used the model framework in Figure 7A-B to calculate the fraction of active cells and mean transcription rate of these active cells. We then compared these simulated population-level responses to normalized dox titration data (Figure 2) to fit binding parameters, the *k_A_* function, and the shapes of the log-normal distributions for *ω_nuc_* and the *EC*_50_ of *k_A_* (Figure 7C; parameter distributions and values in Figure S7D and Supplementary Table 2).

The model successfully reproduced the graded responses in the fraction of active cells and mean expression level of active cells (Figure 7C). These graded population-level responses were produced by switch-like activation responses (effective Hill exponent ∼24, essentially a perfect switch) at the single-cell level (Figure 7D), where the switching point varied between cells due to underlying heterogeneity in nucleosome binding and synTF activation strengths. To examine the predicted molecular states driving these transcriptional responses, we simulated a population of cells with a synTF input level that activated 50% of cells, computationally separated active and silent cells, and compared predicted synTF binding distributions between these groups (Figure 7E). The model predicted synTF binding distributions that were consistent with our experiments: substantial synTF binding was predicted for both On and Off cells, with Off cells having a slightly left-shifted binding distribution compared to the On cells (Figure 7E; compare to Figure 6F).

We then investigated the gene expression dynamics predicted by this model by stochastically simulating the model in Figure 7A using the Gillespie algorithm^77^ for a population of cells with the fit parameters. We set an intermediate synTF input level at which only a fraction of the cells in the population activated, generating a bimodal gene expression distribution (Figure 7F, right). We then performed an *in silico* sorting experiment by separating high-expression and low-expression cells at a certain timepoint. As expected, predicted gene expression states following sorting were stable at the single-cell level, similar to our experimental observations (Figure 7F, left; Figure 3E). In addition, we performed an *in silico* dox upshift and downshift experiment. During the transient dox upshift, we observed that all low-expression cells transitioned to the high-expression state. Following the return to the lower dox level, these cells returned to the low-expression state while the originally high-expressing cells remained in the high-expression state (Figure 7F). These results closely resemble our experimental results (Figure 4H; Figure 7F).

One additional assumption in this modeling framework is that bimodal gene expression distributions are produced primarily by variability in the activation rate *k_A_*. This would predict that increasing synTF input at a reporter gene with a basally active promoter, such that all cells start in the active state, would produce unimodal shifts in gene expression in the cell population. To test this prediction, we recruited rTetR-VP48 in K-562 cells containing an AAVS1-integrated reporter gene with the minCMV promoter replaced with the constitutively active PGK promoter^15^. Consistent with this modeling framework, we observed a unimodal increase in Citrine expression in the cell population as dox was increased (Figure 7G).

Together, these results indicate that incorporating chromatin-mediated heterogeneity in synTF binding and activation strengths into a two-step model of gene expression, with synTF binding separately controlling the probability of a cell being active and the mean expression level of active cells, is sufficient to recapitulate all observed features of transcriptional response functions in the Tet-On system, both at the population and single-cell level. While here we made a number of simplifying assumptions and modeling design choices in our kinetic and thermodynamic models and it is likely many other frameworks could describe our data as well, the qualitative features used by the model to explain transcriptional responses are fairly simple and general: synTF input separately regulates the fraction of active cells and their mean expression level, and locus-encoded variability between cells results in a distribution of switch-like, monostable single-cell response functions across the population. This distribution produces graded fractional responses to increasing synTF input at the population level, generating bimodality at intermediate input levels, and stable all-or-none responses at the single-cell level. This framework also predicts that promoters with constitutive basal levels of transcription should yield unimodal, rather than bimodal, responses in the population, and this prediction was borne out by experiments.

### Activation domains can decouple the fraction of active cells and the mean expression level of active cells in the Tet-On system

In the modeling framework presented in Figure 7A-B, we assumed that the amount of synTF bound at the reporter gene determines both the probability of a cell being active and the mean expression level of active cells. If this were the case, the relationship between the fraction of active cells and their mean expression level should be invariant across different methods of perturbing synTF input (i.e., the same level of synTF binding should always produce the same fraction of active cells and mean expression level of active cells whether that level of binding is produced by treating cells with a certain dox concentration or by changing the number of binding sites in the reporter gene). To test this prediction, we compared the fraction of active cells and the mean expression level of active cells across several of our experiments: an experiment varying dox with a K-562 reporter line containing 9 TetO sites, an experiment varying dox with a K-562 reporter line containing 5 TetO sites, and an experiment with constant dox but in K-562 reporter lines containing 0 to 9 TetO sites. We observed that, for all of these experiments, the relationship between the fraction of active cells and their mean expression level collapsed to a single curve (Figure 7H). This suggests that the relationship between these variables is fixed regardless of the specific method used to perturb synTF input.

The specific relationship between these two variables could be explained by the VP48 activation domain differentially interacting with multiple transcriptional cofactors that affect transcription through distinct mechanisms (e.g. interactions with the chromatin remodelers could primarily affect the probability that a cell is active while direct interactions with general TFs or polymerase could primarily affect the transcription rate). We therefore wondered if this relationship was specific to VP48 or more fundamental to gene expression in the Tet-On system.

To explore this concept, we re-analyzed previously collected flow cytometry time course data^16^ for 10 other human activation domains fused to rTetR and recruited with high dox (1000 ng/mL) in K-562 cells containing the same reporter gene described above, then quantified the fraction of active cells and their mean expression levels after 5 days of recruitment (Figure 7I; Figure S7E-F). Interestingly, most tested activation domains deviated from the fraction on vs. mean expression level relationship measured for VP48 (Figure 7I). Activation domains could have similar fractions of active cells but vary widely in the expression level of these active cells, and vice versa. These results show that activation domains can decouple the fraction of active cells from their mean expression level, possibly through domain-specific interactions and interaction affinities with transcriptional cofactors that primarily work on separate steps of the transcriptional activation process.

Of particular interest, we identified a handful of activation domains that resulted in activation of close to 100% of the cells in the population, but with mean expression levels that varied over an order of magnitude (Figure 7J). These synTFs could prove useful in applications that require uniform, inducible control of transgene expression over a range of gene expression setpoints. This would allow for the study of downstream biological processes following the induction of a particular protein at different doses without the complications of bimodality.

## Discussion

Developing a quantitative understanding of the relationship between transcription factor input and gene expression output is a central goal in the field of gene regulation. Here, we have combined single-cell and single-molecule approaches to better understand transcriptional responses in a minimal synthetic expression system in human cells. We observed that transcriptional responses are graded at the population level but are largely all-or-none at the single-cell level, and that synTF input regulates both the fraction of active cells and the mean expression level of these active cells (Figures 1-2). This fractional activation at intermediate synTF levels is primarily caused by cell-to-cell variation in highly ultrasensitive transcriptional response functions that are stable over time (Figure 3-4). We found that this variation was encoded in the local environment of individual reporter genes and is associated with both reduced synTF binding and weaker activation when bound (Figures 5-6). We then developed a mathematical model of gene regulation incorporating chromatin-mediated heterogeneity that accounted for all of our experimental observations and that correctly predicted that promoters with basal levels of transcription should yield unimodal response functions across the population. Finally, we demonstrated that synTF activation domains can decouple the fraction of active cells from the mean expression level of active cells (Figure 7).

Graded population level and all-or-none single-cell transcriptional responses have been observed in many, but not all^78,79^, studies of synthetic expression systems in mammalian cells^7,14,16–21,26,80–83^. These studies typically used lentivirus or piggyBac transposase to randomly introduce reporter genes into cells. The resulting heterogeneity in reporter number and integration site likely contributes to the observed heterogeneity in transcriptional response functions. They also typically have not quantified TF expression levels in individual cells, which could also contribute to heterogeneous responses. Here, we have built upon these studies and demonstrated that many previously observed characteristics of transcriptional response functions hold even when reducing these sources of variability. These features of transcriptional response functions therefore seem to be a more general aspect of mammalian synthetic expression systems.

Our investigation supports a framework where TFs regulate two separable steps of transcriptional activation: turning a gene from a silent to an active state, then determining the rate of transcription once active. This framework is conceptually similar to previously proposed models of transcriptional regulation in eukaryotes where TFs regulate mechanistically distinct processes involved in gene expression. These include seminal works in yeast that demonstrated nucleosome positioning and occupancy can decouple the activation threshold from its dynamic range^29,84^, studies of transcriptional activation in *Drosophila* development that proposed TFs actively regulate transitions from silent to active chromatin states in addition to regulating polymerase recruitment once active^8,9,36^, and numerous models of transcriptional cycles^31,32^ and transcriptional bursting. It is likely that transcriptional dynamics in the active state measured here are well described by kinetic cycling or transcriptional bursting models. However, our results suggest that there is an additional layer to transcriptional regulation where TFs interface with (sometimes heterogeneous) local chromatin environments to mediate switch-like transitions from an inactive to an active gene expression state prior to polymerase recruitment, and that this layer is crucial for understanding single-cell and population-level transcriptional responses in mammalian cells.

Our mathematical model of transcriptional regulation in the Tet-On system inferred that single-cell transitions from the silent to active state are switch-like; the fitted Hill exponents for the single-cell responses were ∼24, essentially a perfect switch. It is well established that chromatin regulation involves reader/writer enzymes that can introduce positive feedback into transcriptional regulation^47,74^, which in principle can generate switch-like responses through bistability. Positive feedback has been previously proposed to explain bimodal gene expression in mammalian synthetic expression systems^26^, and other models using positive feedback have been developed to describe transcriptional regulation via enhancer-promoter contacts, including through the nucleation of transcriptional condensates^27^. However, we detected almost no hysteresis, indicating a lack of bistability, and instead found largely monostable steady state transcriptional response functions. We suspect that biochemical feedback in chromatin remodeling plays an important role in generating the observed high levels of ultrasensitivity, but does so without generating bistability^85^. Energy expenditure in gene regulation is also likely related to the switch-like nature of the responses, as we were unable to model such high sensitivities with an equilibrium framework^10,34,65,67,86–89^. Future work connecting molecular mechanisms to this ultrasensitivity will be important for dissecting the role of energy expenditure and feedback in generating switch-like transcriptional responses that are largely monostable.

Despite most (∼95%) cells in the population containing the reporter gene in the same genomic locus, we found that the observed variability in monostable transcriptional response functions is primarily encoded in the local environment of the reporter gene rather than through a cell-level variable (e.g., cofactor expression levels). Our data support a model where individual reporter genes exist in a spectrum of chromatin states that produces a distribution of activation potentials across the cell population. This variation in activation potential, which is heritable and persists through periods of high induction and activation, results in a fraction of cells not responding to synTF input at lower induction levels. Epigenetic chromatin states could both be heritable and produce an effectively continuous distribution of states in the cell population. In addition, it is known that gene expression states and DNA accessibility are associated with certain chromatin states^12,70–73^, and epigenetic marks have been found to affect features of transcriptional responses^13,90–92^ and the bimodal gene expression distributions of pluripotency markers in embryonic stem cells^93^.

Our footprinting data showed that the cells that failed to activate at low induction levels are not completely incapable of binding the synTF, but instead have reduced synTF binding and increased nucleosome occupancy. Interestingly, these data also revealed that levels of synTF binding were not solely predictive of gene expression state and level: cell populations with the same average synTF binding could belong to sorted On or Off conditions. These results suggest that there is not a simple map between the number of bound TFs and active gene expression even in simple synthetic contexts and that instead chromatin can modulate and decouple this relationship.

Identifying the relationship between specific chromatin states and their effects on distinct features of transcriptional response functions, particularly the mechanisms by which they reduce synTF binding and activation strength when bound, will be important for further developing a quantitative understanding of mammalian gene regulation across diverse contexts and genomic loci beyond AAVS1. While here we analyzed a synthetic expression system integrated in this particular locus, it is likely that the principles connecting chromatin states to transcriptional responsiveness are fundamental and could be related to endogenous contexts where bimodal gene expression distributions and all-or-none responses are observed, such as for random monoallelic expression^94^. In addition, it will be interesting to investigate where in the cell-engineering process this chromatin heterogeneity arises in the cell population and to see if it can be minimized through different engineering strategies.

Finally, we found that the relationship between the probability of a cell being active and its mean expression level once active is specific to individual activation domains (i.e., some activation domains efficiently turn cells on, but to a low expression level, while others activate fewer cells but to a higher level). Transcriptional activation domains interact with distinct cofactors that are thought to function in distinct steps of gene expression, such as chromatin remodeling, transcriptional initiation, pausing, and elongation^31,32,35,44,95^. We speculate that the differential effects activation domains exert on gene regulation in our system are caused by differential interaction affinities with cofactors that efficiently turn genes on versus ones that drive high levels of expression at active genes. Mapping these interactions to quantitative features of transcriptional responses will further our predictive understanding of mammalian gene regulation. Practically, identifying these principles could lead to the development of expression systems that drive more uniform gene expression responses across cell populations (such as the synTF characterized here) to complement existing methods for achieving setpoint control in gene regulation^14,40,96,97^. More fundamentally, identifying the functional relationship between chromatin states, transcriptional cofactors, and both single-cell and population-level features of transcriptional response functions will advance our basic understanding of mammalian gene regulation that can be generalized across contexts.

## Resource Availability

Raw and processed next generation sequencing data from single-molecule footprinting are available through the Gene Expression Omnibus (GSE345733). All reporter plasmids and synTFs used in this study will be made available on Addgene or upon request. Source data locations, plasmid maps, flow cytometry data, flow cytometry processing code, flow cytometry analysis code, and code used for mathematical modeling have been uploaded to Zenodo (DOI: 10.5281/zenodo.22308911). Requests for further information and resources should be directed to and will be fulfilled by the lead contact, Lacramioara Bintu.

## Acknowledgments

The authors would like to thank all members of the Bintu and Ferrell labs, as well as Gavin Schlissel, Martha Cyert, Jonas Cremer, and especially Alistair Boettiger, for helpful discussions and valuable feedback. This work was supported by the NIH T32GM007276 (E.J.C.), Stanford H&S Graduate Research Opportunity Fund (E.J.C.), NIH T32GM141828-01A1 (C.J.A.), NSF GRFP DGE-2146755 (C.R.M.), Stanford VPGE EDGE Fellowship (C.R.M.), NIH R35GM128947 (L.B.), and NIH R35GM131792 (J.E.F.).

## Author Contributions

Conceptualization: E.J.C., L.B., and J.E.F.; data curation: E.J.C.; formal analysis: E.J.C.; Investigation: E.J.C., C.R.M., and C.J.A.; methodology: E.J.C.; software: E.J.C. and C.R.M.; visualization: E.J.C.; writing - original draft: E.J.C.; writing - reviewing and editing: E.J.C., C.R.M., C.J.A., L.B., and J.E.F.; project administration: L.B. and J.E.F.; resources: L.B.; funding acquisition: E.J.C., C.R.M., L.B., and J.E.F.; supervision: L.B. and J.E.F.

