## Supplemental Information for "Stable epigenetic states set single-cell activation thresholds in mammalian expression systems"

**This PDF includes:**

Supplemental Figures 1-7

Methods

References for Supplemental Materials

### Supplemental Figures

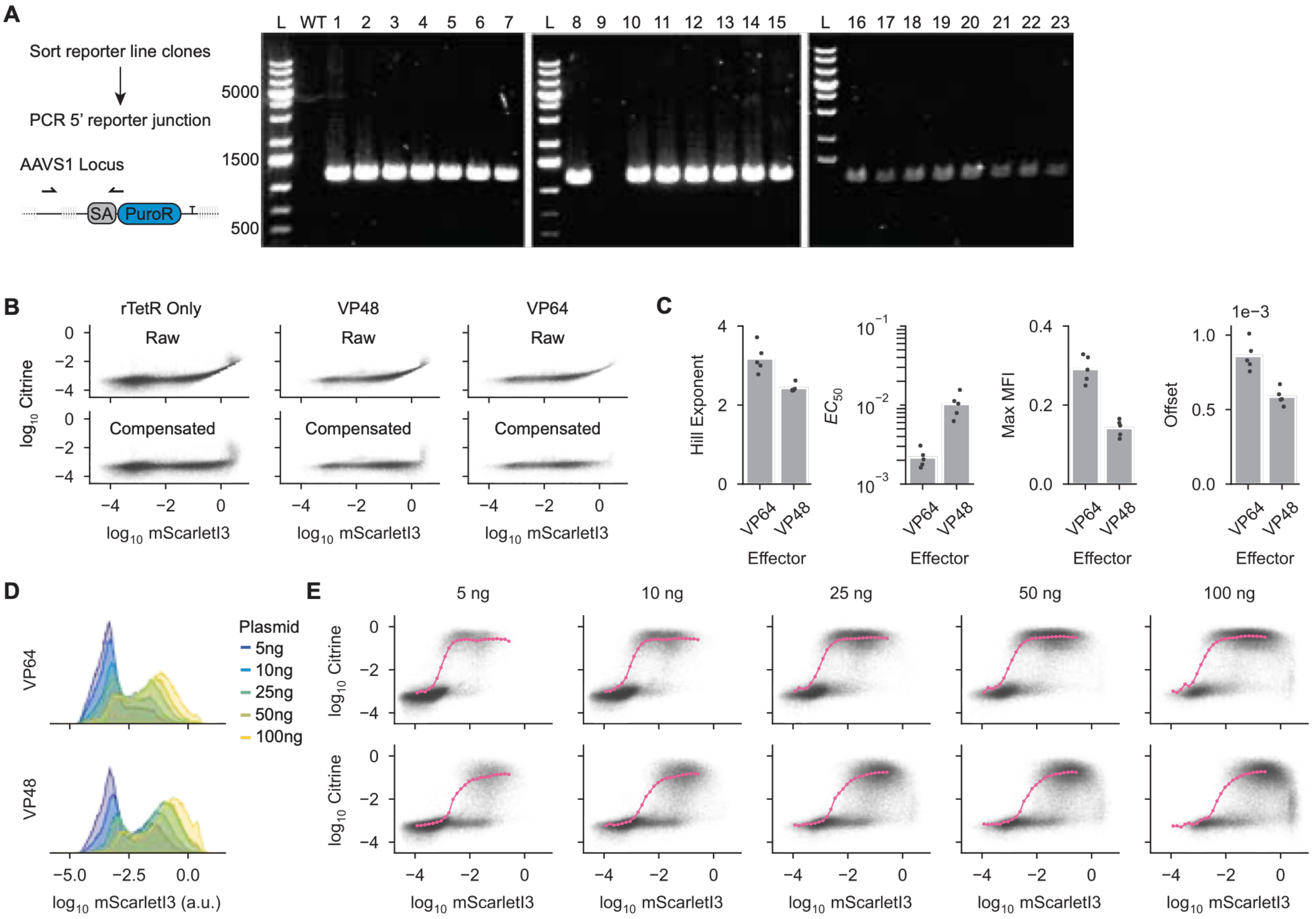


**Figure S1:** Characterization of reporter integrations and response functions measured with transient transfections.

A) Strategy for determining on-target reporter integration rate. K-562 cells were electroporated with TALEN expression plasmids and the reporter plasmid, then selected with puromycin. Single clones were sorted for synTF expression and expanded. After genomic DNA extraction, the 5' AAVS1-reporter junction was amplified using PCR. The expected PCR band is only present in clones with successful integration of the reporter in the AAVS1 locus. Numbers indicate clones. WT = no reporter cells. L = ladder.

B) Flow cytometry measurements for control transfection of rTetR-mScarletI3 into HEK-293T reporter cells, before and after compensating mScarletI3 bleed-through into the Citrine channel, for rTetR only, rTetR-VP48 with no dox, and rTetR-VP64 with no dox. A linear function was used to model bleed-through.

C) Fit parameters describing the Hill Functions shown in Figure 1D. Individual points represent different total amounts of plasmid transfected. An offset reflecting different signal baselines in the Citrine channel was included in the fit, and the difference in the fit offset values are subtracted from the VP64 measurements in Figure 1.

D) mScarletI3 (synTF) expression distributions in HEK-293T reporter cells with increasing amounts of total plasmid transfected for VP64 (top) and VP48 (bottom).

E) Flow cytometry measurements after transfecting varying amounts of VP64 (top) or VP48 (bottom) into reporter cells. 1000 ng/mL dox was added 24 hours after transfection, and measurements were taken 24 hours after dox addition. Pink lines represent the mean Citrine expression in cells binned by mScarletI3 level, as in Figure 1D.

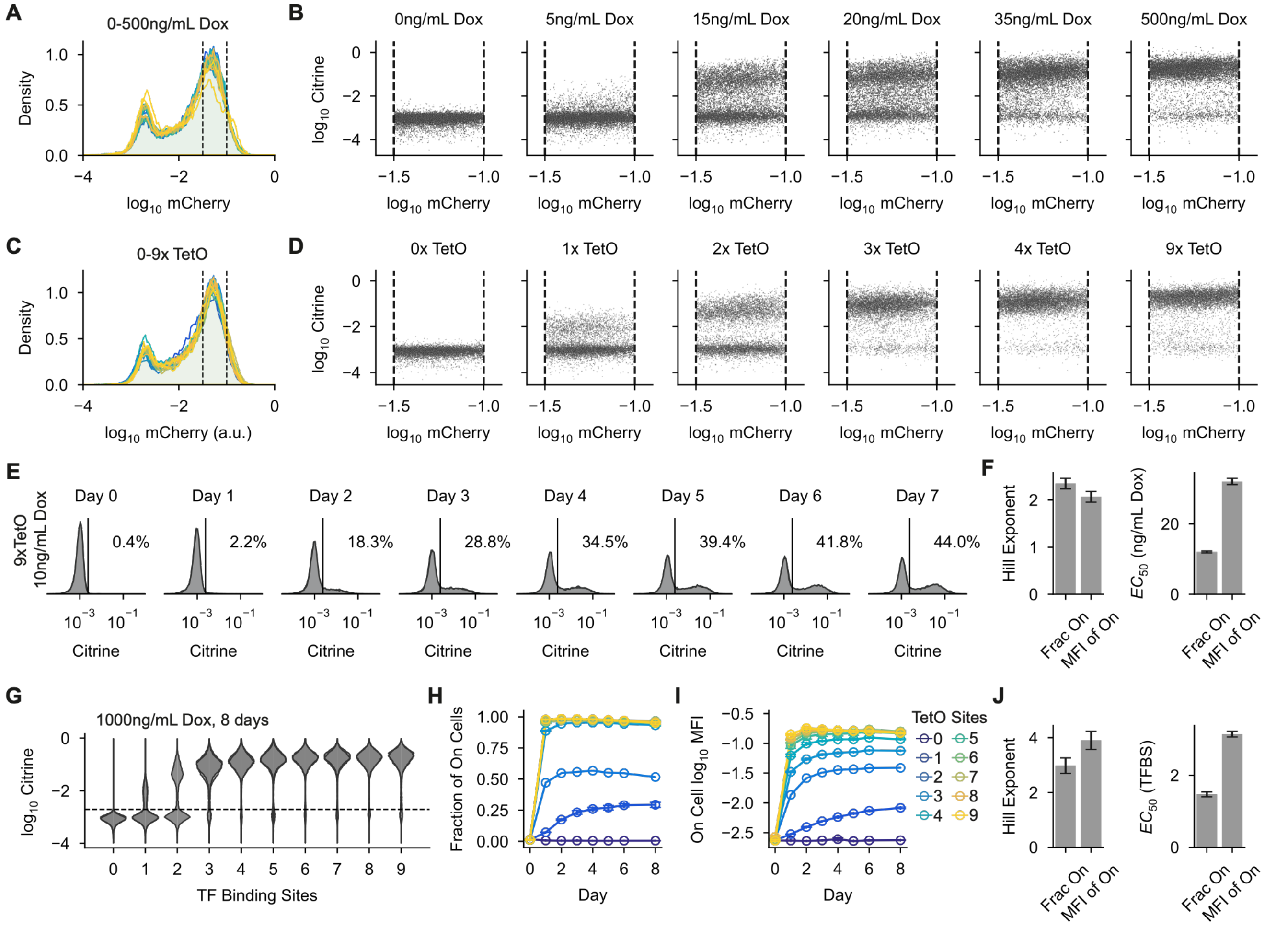


**Figure S2:** Validation of using dox concentration and number of synTF binding sites to measure transcriptional response functions over long timescales in K-562 cell lines stably expressing rTetR-VP48.

A) mCherry expression distributions in K-562 cells containing a reporter gene with 9 TetO sites. Measurements were taken after 7 days of synTF recruitment. Colors represent different dox concentrations added to the cells; dashed lines represent the upper and lower bounds used to gate cells with similar synTF expression levels.

B) Flow cytometry measurements of the gated cells in A for several dox concentrations. Dashed lines represent the mCherry gate from A.

C) mCherry expression distributions in K-562 cell lines containing reporter genes with 0 to 9 TetO sites. Measurements were taken after 8 days of synTF recruitment using 1000 ng/mL dox. Colors represent different numbers of binding sites; dashed lines represent the upper and lower bounds used to gate cells with similar synTF expression levels.

D) Flow cytometry measurements of the gated cells in C for several numbers of synTF binding sites. Dashed lines represent the mCherry gate from C.

E) Citrine expression distributions over time during synTF recruitment at 10 ng/mL dox in K-562 cells containing a reporter gene with 9 binding sites.

F) Parameter fits for the Hill functions shown in Figure 2E. Error bars represent the SD of the fit for each parameter.

A) Citrine expression distributions following 8 days of synTF recruitment in cells containing reporter genes with 0 to 9 TetO sites at 1000 ng/mL dox. Two biological replicates are shown. The dashed line represents the gate used to call active cells in H-I.

B) Fraction of active cells over time for the conditions in G. Error is the SD of two biological replicates.

C) Mean expression level of active cells over time for the conditions in G. Error is the SD of two biological replicates.

D) Parameter fits for the Hill functions shown in Figure 2F. Error bars represent the SD of the fit for each parameter.

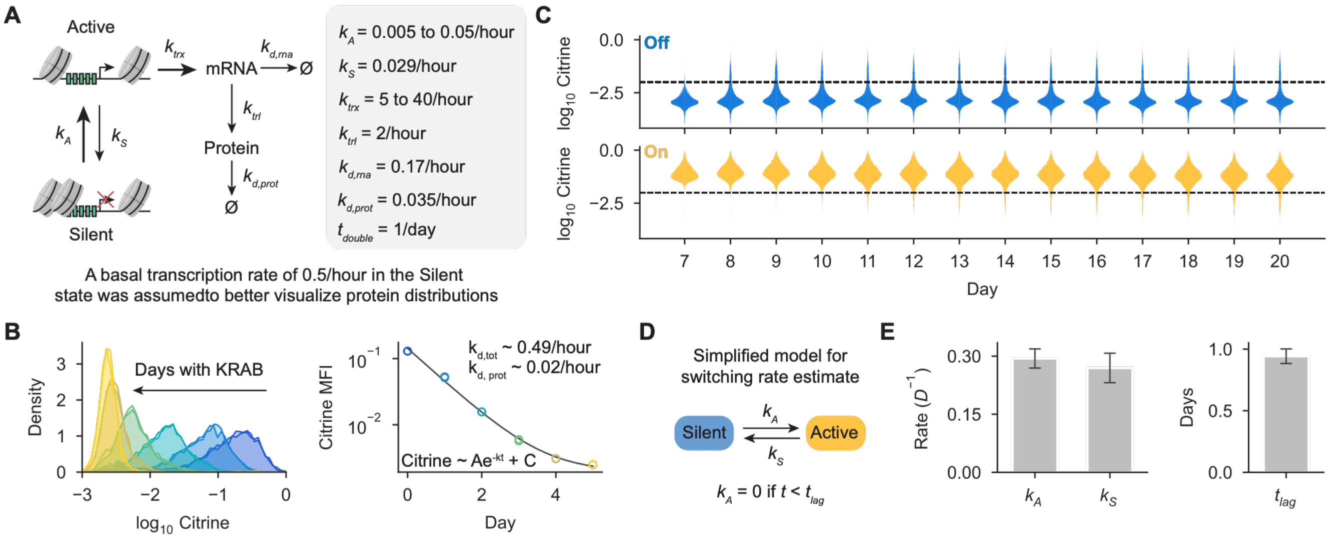


**Figure S3:** Analysis of two-state bursting model and gene expression state stability in the Tet-On system.

A) Parameter values used to simulate single-cell gene expression dynamics in Figure 3A-C. For simplicity, cells were assumed to divide every 24 hours, but the time of first division was random to un-synchronize the cells. A basal transcription rate of 0.5/hour was assumed to occur in the Silent state to better visualize the Off population of cells at the protein level (see Methods).

B) Citrine expression distributions while recruiting the strong transcriptional repressor rTetR-KRAB;ZNF10 in K-562 cells containing a reporter gene with 9 TetO sites upstream of the constitutively active pEF promoter using 1000 ng/mL Dox. Darkest blue = day 0 of recruitment, lightest yellow = day 5 of recruitment. An exponential decay model fit to the mean Citrine expression on each day is shown on the right. From this fit, the Citrine degradation rate was estimated assuming a cell-doubling time of 24 hours.

C) Complete Citrine expression distributions corresponding to the line plots in Figure 3E for sorted On and Off cells. Two biological replicates are shown. Dashed lines represent the gate used to call On and Off fractions in Figure 3E.

D) Simplified two-state model used to estimate gene expression switching rates from the data in Figure 3E. Protein turnover was assumed to be fast enough that the fraction of active cells roughly corresponded to the fraction of cells in the Active state. To better estimate switching rates and account for the time it took for enough protein to accumulate for a cell to be called as On, a lag time was included in the model (see Methods).

E) Parameter fits for the model in D, used to estimate population re-equilibration rates under the dynamic equilibrium model. Error bars represent the SD of each fit parameter. The model was fit to days 0 to 7 of the recruitment time course in Figure 3E.

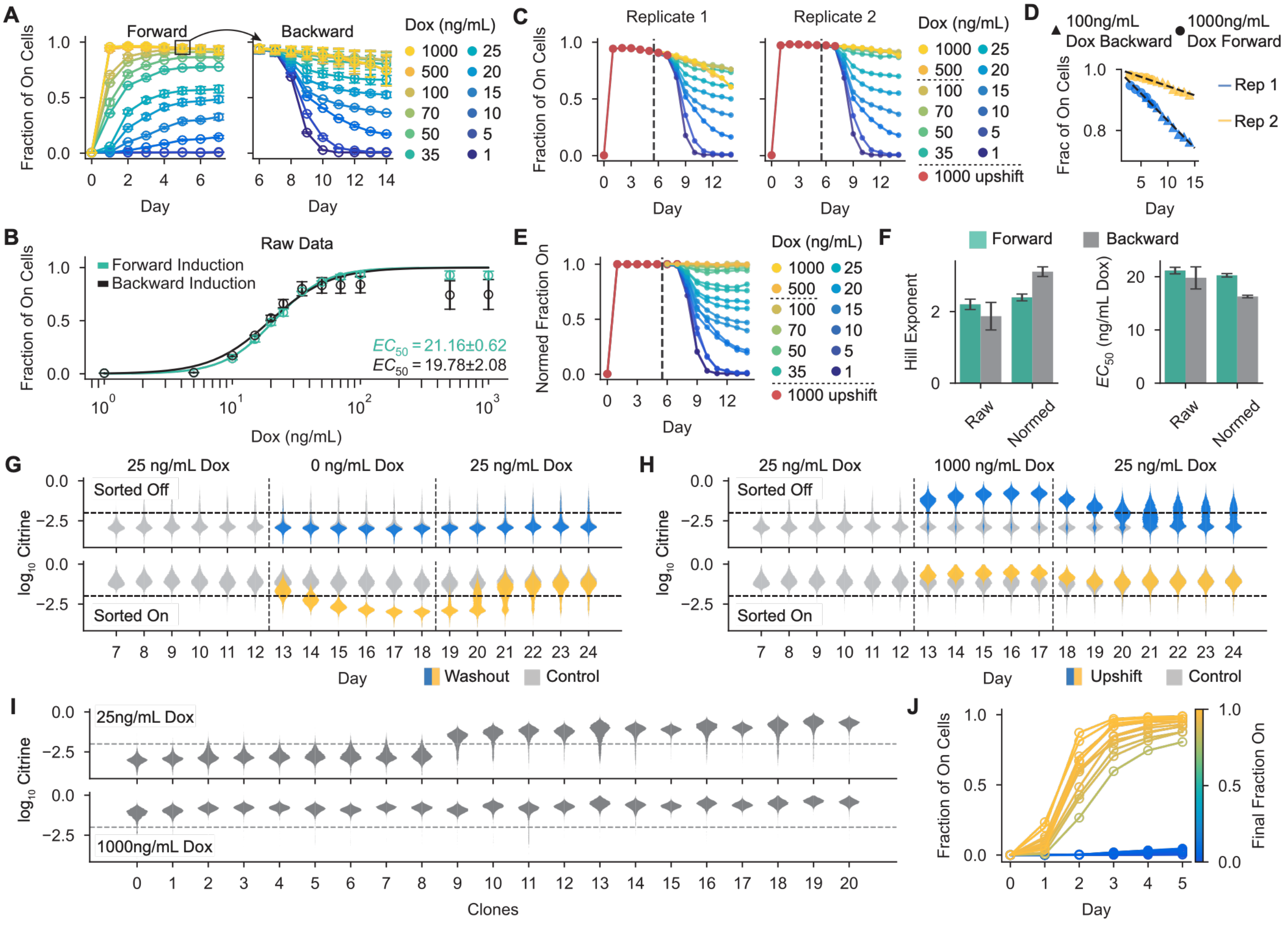


**Figure S4:** Normalization of forward and backward inductions in the Tet-On system, analysis of hysteresis in sorted populations, and rTetR-VP48 recruitment in clonal cell lines.

A) Un-normalized fraction of active cells over time for the forward and backward recruitment time courses shown in Figure 4D. Error represents the SD of two biological replicates; colors represent different dox concentrations.

B) Final fraction of active cells for the un-normalized forward and backward inductions, as in Figure 4E.

C) Un-normalized fraction of active cells over time from A, but separated by replicate. We observed that the rate of background silencing accelerated for 500 ng/mL dox and 1000 ng/mL dox, while it seemed constant and the same as the pre-downshift 1000 ng/mL dox condition for the lower dox doses. Background silencing was replicate-specific.

D) Linear functions fit to the fraction of active cells for the pre-downshift 1000 ng/mL dox conditions and post-downshift 100 ng/mL dox condition. Fits were used to model replicate-specific background silencing at lower dox doses.

E) Normalized fraction of active cells over time for forward and backward inductions, as in Figure 4D but plotting replicates separated. 500 ng/mL and 1000 ng/mL conditions were assumed to have 100% of cells active without background silencing, 0-100 ng/mL dox conditions were assumed to have a linear rate of silencing determined in D.

F) Parameters from Hill function fits to the normalized and un-normalized forward and backward inductions (Figure 4E and Figure S4B respectively). Error bars represent the SD of each fit parameter.

G) Full Citrine expression distributions for the line plots shown in Figure 4G. Vertical dashed lines mark washout window; horizontal dashed lines represent the On cell gate used for Figure 4G.

H) Full Citrine expression distributions for the line plots shown in Figure 4H. Vertical dashed lines mark upshift window; horizontal dashed lines represent the On cell gate used for Figure 4H.

I) Citrine expression distributions following 5 days of rTetR-VP48 recruitment in clonal K-562 lines containing a reporter gene with 5 TetO sites at 25 ng/mL and 1000 ng/mL dox. Dashed lines represent the On cell gate used in J.

J) Fraction of active cells over time for the clonal cell lines shown in I

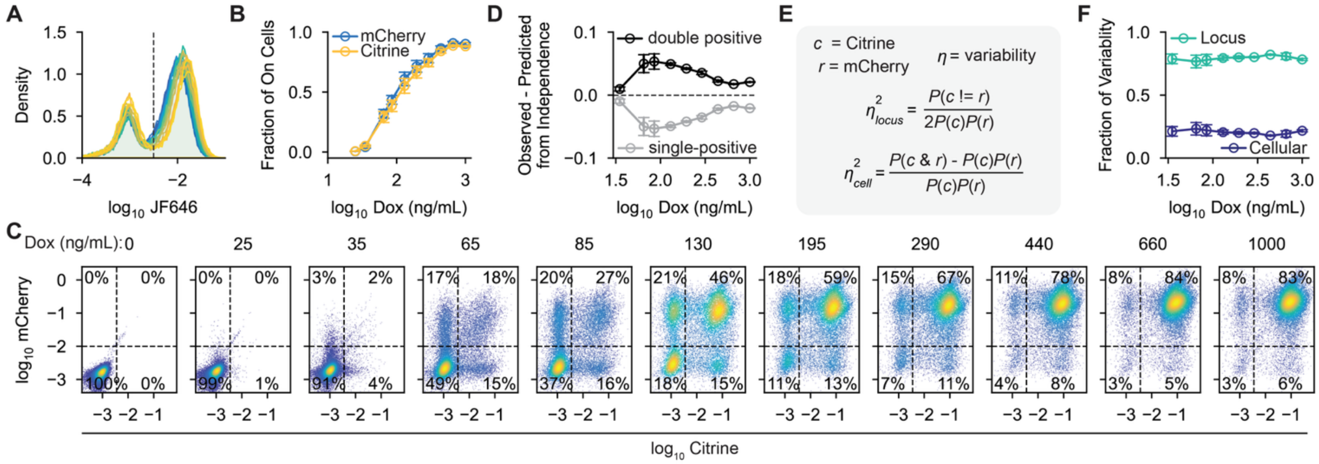


**Figure S5:** Decomposition of variability in transcriptional response functions into cell-level vs locus-level effects.

A) JF-646 fluorescence distributions after 6 days of synTF recruitment for the dual-reporter cell line described in Figure 5. Colors represent different dox concentrations; the dashed line represents the gate used to select cells expressing the synTF for downstream analysis.

B) Fraction of cells expressing Citrine or mCherry after 6 days of synTF recruitment in the dual-reporter cell line. Error represents the SD of two biological replicates.

C) Flow cytometry measurements for Citrine and mCherry reporter expression after 6 days of synTF recruitment, as in Figure 5C but for all dox doses analyzed. Dashed lines represent the gates used to call gene expression states.

D) Deviation of observed fraction of double-positive and single-positive cells from predicted fractions assuming complete independence between the reporters. Error represents the SD of two biological replicates. Deviations were calculated within replicates. The dashed line represents where no deviation would lie.

E) Framework used to calculate the relative contribution of locus-level and cell-level effects on variability between transcriptional response functions. We employed a binary version of the approach taken by Elowitz et al.[1] and made the simplifying assumption that reporters activated symmetrically on average across the population.

F) Fraction of variability from locus-level and cell-level effects, as in Figure 5F but as a function of dox concentration.

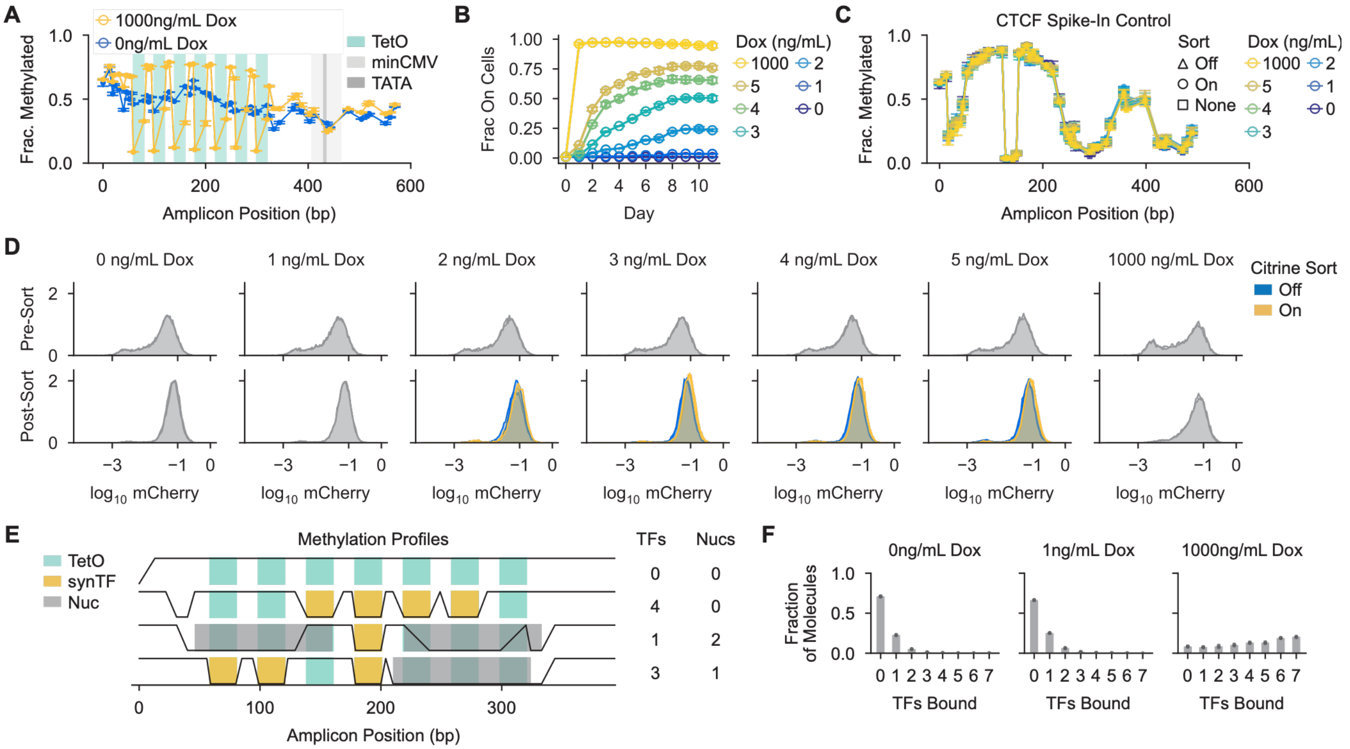


**Figure S6:** Single-molecule footprinting controls and examples of molecular states.

A) Bulk methylation profiles for 1000 ng/mL and 0 ng/mL dox footprinting conditions shown in Figure 6C. Positions of TetO sites, the minCMV promoter, and the TATA box are indicated. Error bars represent the SD of two biological replicates.

B) Pre-sort fraction of active cells over time during synTF recruitment in the SMF reporter line at a range of dox concentrations. Error bars represent the SD of two biological replicates.

C) Bulk methylation profiles for a reporter gene containing a single CTCF binding site spiked into the SMF conditions shown in Figure 6. ~50k CTCF cells were added to each condition immediately before nuclei extraction.

D) mCherry (synTF expression) distributions before (top) and after (bottom) sorting cells for SMF.

E) Example single-molecule methylation profiles and molecular states assigned to them by a probabilistic binding model[2]. TetO site locations are shown in teal, synTF binding events are shaded yellow, and called nucleosomes are shown in grey. For each molecule, unmethylated GpC sites are down while methylated GpC sites are up. Example molecules were taken from cells treated with 5 ng/mL dox and sorted On.

F) SynTF binding distributions for cells treated with 0, 1, and 1000 ng/mL dox, calculated as described in Figure 6F. Black points indicate biological replicates for each condition. Binding distributions were calculated using a maximum likelihood model, as previously described[2].

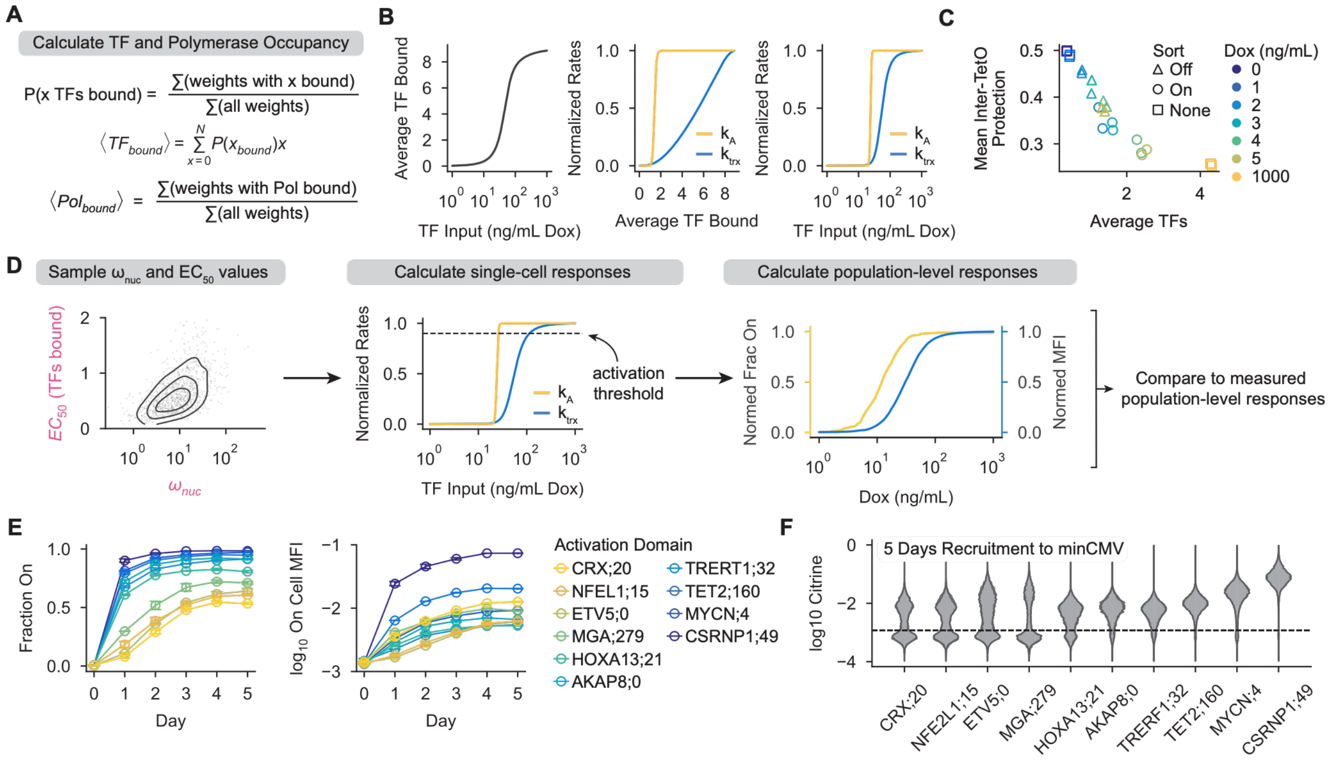


**Figure S7:** Detailed description of combined thermodynamic and kinetic model incorporating local variability to describe gene expression in the Tet-On system.

A) Method used to calculate kinetic rates from the thermodynamic model described in Figure 7B. Briefly, the statistical weights of all enumerated molecular states are used to calculate the partition function of the system, which is then used to calculate average synTF and polymerase occupancy. These average occupancies are then used to calculate the activation rate and the transcription rate for the kinetic model in Figure 7A.

B) Relationship between synTF input, in units of dox, and the average number of synTFs bound and the normalized kinetic rates of the system. Relationships are shown for a single example cell that represents the mean of the distribution using the parameters in D.

C) Relationship between average synTF occupancy and the mean inter-TetO protection, taken as a proxy for average nucleosome occupancy.

D) Process used for fitting the model in Figure 7A-B while incorporating cell-to-cell variability in nucleosome binding and activation sensitivity. Pairs of parameters were sampled for 300 cells (left), then used to calculate normalized rates of activation and transcription in single cells (middle). The fraction of cells with an activation rate above 0.9 and mean transcription rate of active cells were then calculated for several synTF input levels and fitted to the normalized fraction of active cells and MFI of active cells shown in Figure 2E. See Methods for details.

E) Fraction of cells expressing Citrine and mean expression level of active cells during recruitment of various synTFs with 1000 ng/mL dox in K-562 reporter cells containing a reporter gene with 9 TetO sites. Error bars represent the SD of two biological replicates.

F) Citrine expression distributions following recruitment of various synTFs with 1000 ng/mL dox for 5 days in K-562 reporter cells containing a reporter gene with 9 TetO sites. Activation domains are referred to by the name of the TF they originate from and the “tile” number they correspond to in a previously published high throughput screen[3] (e.g. CRX;20 indicates amino acids 201 to 280 from the human TF CRX). The dashed line represents the gate used to call On cells in E.

### Methods

*Plasmid Cloning*

SynTFs were cloned using effector domains ordered as gBlocks from IDT and assembled into the backbones pJT126 (Addgene no. 161926), backbone EC063 (Figure 1), or backbone EC051 (Figure 5) using GoldenGate cloning. Golden Gate assembly was performed in 10 μL reactions using 80ng of plasmid backbone, 5ng of insert (2:1 molar ratio of insert/backbone), 1 μL of 10x T4 DNA Ligase Buffer (NEB, B0202S), 0.25 μL T4 DNA Ligase 2,000,000U/mL (NEB, M0202T), 0.75 μL Esp3I restriction enzyme (NEB, R0734L), and nuclease-free ddH2O. Typically, 35 rounds of digestion-assembly cycles were performed in a thermocycler (Bio-Rad).

To construct backbone plasmids, pJT126 was modified using Gibson assembly. Reporter plasmids with different numbers of binding sites (plasmids EC040-EC048) were designed by sequentially scrambling the sequences of TetO sites from the 5' end of the 9xTetO minCMV reporter plasmid DY032 (Addgene no. 161928). These new binding site arrays were ordered in two halves split in the middle of the binding region as gene fragments from TWIST and assembled into backbone CL056 using three-part Gibson assembly. The mCherry reporter (EC057) used for the multi-output experiments was engineered by replacing the Citrine in plasmid EC044 using Gibson assembly.

For all Gibson assemblies, vectors were obtained using restriction enzyme digestion of the relevant backbone followed by gel extraction (Zymo, D4001T). Insert fragments were obtained using PCR from plasmid templates using Q5 Hot Start High-Fidelity 2X Master Mix (NEB, M0494S) followed by gel purification or ordered as gene fragments from IDT or TWIST. 20-30bp of overhangs were used for all assemblies. Reactions were performed in 20 μL total with 50-100ng of gel extracted vector, a 2-fold molar excess of insert DNA (resuspended gene fragments or PCR product), and 10 μL of NEBuilder HiFi DNA Assembly Master Mix (NEB, E2621L). Reactions were incubated at 50C for 60 minutes, then 2 μL of assembly product was transformed into Mix and Go chemically competent cells. All cloned constructs were verified using whole-plasmid sequencing (Plasmidsaurus).

All plasmids, oligos, and activation domain sequences used in this study are listed in Supplementary Table 1.

*Cell Culture and Engineering*

All experiments presented here were performed in K-562 cells (ATCC, CCL-243, female) except for those in Figure 1. Cells were cultured in a controlled humidified incubator at 37C and 5% CO2, in RPMI 1640 (Gibco, 11-875-119) media supplemented with 10% fetal bovine serum (FBS) (ThermoFisher, A3382101) and 1% penicillin-streptomycin-glutamine (Gibco, 10-378-016). Cells were kept at a confluency of less than 1 million cells/mL of culture. HEK-293T-LentiX (Takara Bio, 632180, female) cells, used to produce lentivirus (described below), were grown in DMEM (Gibco, 10569069) media supplemented with 10% FBS (ThermoFisher, A3382101) and 1% penicillin streptomycin (Gibco, 15-140-122).

To engineer HEK-293T reporter cells, ~50k cells were plated in a 6-well tissue culture plate and grown overnight. Then, 500ng of DY032 reporter plasmid (Addgene no. 161928) was co-transfected with 250 ng each of TALEN-L (Addgene no. 35431) and TALEN-R (Addgene no. 35432). Transfections were performed using 5 μL Lipofectamine LTX and 2.5 μL PLUS reagent (ThermoFisher, 15338030) in 250 μL of Opti-MEM (Gibco, 31-985-070). After 48 hours, the cells were treated with 1000 ng/mL puromycin (Gibco, A1113803) until all non-resistant cells were killed and a population where the reporter was stably integrated in the intended locus remained (5-7 days).

K-562 reporter cell line generation was performed as previously described for cell lines containing a single reporter gene[4]. Briefly, DY032 (9xTetO minCMV), JT039 (9xTetO pEF, Addgene no. 161927), JS082[2] (7xTetO minCMV for SMF), CL026 (9xTetO PGK, Addgene no. 196545), and EC040-EC048 (0-8xTetO) Citrine reporter cell lines were created by electroporating 1 million K-562 cells with 1000 ng of reporter donor plasmid and 500 ng of each TALEN-L and TALEN-R plasmid using program T-016 on the Nucleofector 2b (Lonza, AAB-1001). After 48 hours, the cells were treated with 500 ng/mL puromycin until all non-resistant cells were killed and a population where the donor was stably integrated in the intended locus remained (5-7 days).

SynTF expression constructs were stably integrated into K-562 cells using lentivirus as previously described. Briefly, HEK-293T LentiX cells were plated in 6-well tissue culture plates in 2mL of DMEM such that they would be at ~80% confluency the next day, grown overnight, and transfected with 750ng of an equimolar mixture of the three third generation packaging plasmids (pMD2.G: Addgene no. 12259; pRSV-Rev: Addgene no. 12253; pMDLg/pRRE: Addgene no. 12251, all gifts from Didier Trono) and 750ng of recruiter construct plasmid using polyethylenimine (Polysciences no. 23966). After 72 hours of incubation, lentivirus was harvested and filtered through a 0.45 μm polyvinylidene fluoride filter (Millipore). Reporter K-562 cells were transduced with the lentivirus by spinfection for 2 hours using 1mL of harvested virus per 300k cells or the equivalent amount of concentrated virus (Takara Bio, 631232). After 48 hours, infected cells were selected with 10 μg/mL of blasticidin (Gibco) until at least 80% of cells expressed the recruiter construct, as measured by mCherry expression via flow cytometry (Bio-Rad ZE5), or cells doubled every 24 hours under selection (~7-10 days).

To generate the dual-reporter cell line (Figure 5), 1 million K-562 cells were electroporated with 500ng of each reporter plasmid (EC044 (4xTetO minCMV Citrine) and EC057 (4xTetO minCMV mCherry)) and 500 ng of each TALEN-L and TALEN-R plasmid using program T-016 on the Lonza Nucleofector 2b. After 48 hours, cells were treated with 500 ng/mL puromycin for 7 days. rTetR-HALO-VP48 (EC052) was stably integrated into the reporter lines using lentivirus. 48 hours later, cells were treated with blasticidin until the surviving cells doubled roughly every 24 hours in the presence of the antibiotic, demonstrating resistance to the selection marker. Cells were treated with 100 ng/mL doxycycline (Fisher Scientific) for 72 hours and Citrine+ and mCherry+ cells were enriched by cell sorting using a Sony Cell Sorter SH800S. Cells were expanded for ~7 days after sorting to allow for reporter protein to dilute and degrade out of the culture before experiments.

These cell lines were not authenticated. All cell lines tested negative for mycoplasma. All experiments were performed with at least two biological replicates at the level of reporter line electroporation, unless indicated otherwise.

*Transient Transfections*

To measure transcriptional response functions using transient transfections (Figure 1), ~20k HEK-293T cells containing a 9xTetO reporter gene (DY032) were plated in separate wells of a 24-well tissue culture plate. The following day, varying amounts of VP48 and VP64 synTF expression constructs (EC066, EC067) were combined with 1 μL Lipofectamine LTX and 0.5 μL PLUS reagent in 50 μL of Opti-MEM according to manufacturer instructions and added dropwise to cells. After 24 hours, 0 or 1000 ng/mL doxycycline was added to each well. After 24 hours of dox treatment, cells were washed with PBS then dissociated with Trypsin-EDTA (0.25%) and resuspended in 0.3mL RPMI for analysis. SynTF and reporter gene expression were then measured by flow cytometry (Bio-Rad ZE5, Everest v.2.3-3.0).

*Recruitment Time Courses*

To measure gene expression dynamics during effector domain recruitment, $4x10^5$ K-562 reporter cells stably expressing the recruiter construct were typically plated in 1mL of media in separate wells of a 24-well plate and either treated with doxycycline (Fisher Scientific) or untreated. Time points were measured by flow cytometry analysis (Bio-Rad ZE5, Everest v.2.3-3.0) by removing 200k cells per condition. Dox was assumed to degrade each day, so fresh dox media was added each day of the time course. During recruitment time courses, cells were always split 1:2 into fresh media each day.

For recruitment assays with cell sorting (Sony Cell Sorter SH800S), K-562 reporter cells were grown in larger volumes of media that were necessary to obtain sufficient cells after cell sorting. Throughout time courses, cells were maintained at a confluency of 4-8x$10^5$ cells/mL and split 1:2 each day to remain consistent with our 24-well plate experiments. For sorting, cells were centrifuged at 300g for 5 minutes and then resuspended at a concentration of 10x$10^6$ cells/mL in the appropriate doxycycline media. Collection tubes were prepared with doxycycline media at the same concentration to minimize the amount of time cells spent without dox during sorting. After a sufficient number of cells were sorted (typically 500k - 1000k), collection tubes were spun down at 300g for 5 minutes and cells were replated at 400k cells/mL in doxycycline media.

*Flow Cytometry Analysis*

Data were analyzed using Cytoflow (v.1.1, https://github.com/bpteague/cytoflow) and custom Python scripts. Events were gated for viability and fluorescence measurements were normalized by FSC. For experiments in K-562 cells, mCherry or JF-646 was used as a proxy for synTF expression level and cells were gated on synTF expression for downstream analysis, as shown in Figure S2. For transient transfections, fluorescence bleed-through from the mScarletI3 channel into the Citrine channel was compensated using synTF-matched transfections with no dox. Cells were binned by mScarletI3 expression, then a linear model was fit relating the mean of each bin to the average Citrine expression in that bin. This model was then used to subtract bleed-through from +dox transfections. Gates used to compute the fraction of active cells are shown for each plot. Typically, we fit a Gaussian model to cell lines treated with no dox to model the distribution of Off cells, then set a threshold that was two standard deviations above the mean of the Off peak to label active cells. For dual-reporter and cell sorting experiments, gates were set roughly halfway between On and Off populations in log space.

To normalize the fraction of active cells during our forward and backward inductions (Figure 4; Figure S4), we modeled background silencing of the reporter gene and adjusted the raw fraction of active cells by removing the computed background silenced fraction. We first noted that the apparent rate of background silencing was dox dependent, with higher dox concentrations leading to more silencing. Cells treated with 500 and 1000 ng/mL dox silenced at the same rate, which accelerated over time, and were assumed to have 100% of active cells based on our forward inductions. For 100 ng/mL dox and lower, we noticed that the rate of silencing appeared constant and the same as the rate of silencing for cells treated with 1000 ng/mL dox before the dox downshift. We therefore fit a linear model to the fraction of active cells over time by combining the 1000 ng/mL dox forward induction and 100 ng/mL dox backward induction, then corrected the fraction of active cells for all lower doses (Figure S4). All normalization was performed within biological replicates because the rate of silencing was replicate-specific.

*HALOtag Staining*

HALOtag staining was performed using the Janelia Fluor 646 HaloTag Ligand (Promega #GA1121) as follows: ligand was resuspended in 35.5 µL DMSO to prepare a 200 µM stock solution, then diluted to a 200 nM working solution in warm RPMI. 2x$10^5$ cells were pelleted per sample by spinning at 300g for 5 minutes then resuspended in 200 µL of working ligand solution. Samples were incubated for 15 minutes in a 37°C 5% $CO_2$ incubator, then pelleted and resuspended in 500 µL RPMI for flow cytometry analysis.

*Single Molecule Footprinting*

Single molecule footprinting experiments and analysis were performed using an updated version of a previously described protocol[2]. Briefly, K-562 reporter cells were treated with varying concentrations of dox for 11 days, then sorted for mCherry expression and based on Citrine expression as described above. 0 ng/mL and 1 ng/mL dox conditions were not sorted on Citrine. 1000 ng/mL dox was sorted for Citrine positive cells to remove background silenced cells after the recruitment time course. Cells were plated in the same dox concentration after sorting.

The following day, 1 million K-562 cells for each condition were aliquoted and ~50k cells containing a reporter gene with a single CTCF binding site (EC089) were spiked into each tube. Cells were then spun down for 5 minutes at 300g in 2-ml Eppendorf tubes in a swing-bucket centrifuge pre-chilled to 4 °C, which was used throughout the protocol. The supernatant was aspirated and cells were resuspended in 200 µL ice-cold PBS. After spinning cells down (5 min, 300g, 4 °C), the supernatant was aspirated using a P200 to avoid cell loss. To lyse cells using the Omni-ATAC protocol, cells were then resuspended in 200 µL of fresh ice-cold lysis buffer (10 mM Tris-HCl pH 7.5, 10 mM NaCl, 3 mM $MgCl_2$, 0.1% NP-40, 0.1% Tween-20, 0.01% digitonin) by pipetting up and down three times. This cell lysis reaction was incubated on ice for 3 min. After lysis, 1.2 ml of ice-cold resuspension buffer (10 mM Tris-HCl pH 7.5, 10 mM NaCl, 3 mM $MgCl_2$, 0.1% Tween-20) was added, and the tubes were inverted three times to mix the contents. Nuclei were then immediately spun down (10 min, 500g, 4 °C). Supernatant was aspirated first with a P1000 and then with a P200 to avoid nuclei loss. Each methylation reaction was thoroughly mixed on ice using 100 µL of cell methylation buffer (10% volume 10× M.CviPI reaction buffer (NEB, B0227S), 300 mM sucrose, 2.13 mM *S*-adenosylmethionine) and 50 µL of 4,000 U $ml^{-1}$ M.CviPI GpC methyltransferase (NEB, M0227L). Each methylation reaction was then incubated at 37 °C with shaking at 1,000 r.p.m. for 7.5 min and quenched with 150 µL gDNA cell lysis buffer, 3 µL RNase A, and 1 µL ProtK all from the Monarch gDNA extraction kit (NEB, T3010). gDNA was extracted following the protocol from the extraction kit, with the exception that 600 µL of gDNA binding buffer was added to 400 µL quenched reaction and added to the extraction column on two spins of 2 min at 1,000g. DNA was quantified using Qubit 1X dsDNA HS Assay Kit on a Qubit Flex Fluorometer. The appropriate dox concentration was added to each buffer used before quenching the methylation reaction.

Methylated gDNA was digested using XbaI and AccI (NEB) to produce a fragment containing the entire intact reporter sequence, and a two-sided SPRI (0.4x followed by 1.8x) was performed to enrich these fragments. DNA was converted using the Enzymatic Methyl-seq Conversion Module (NEB) following the manufacturer's instructions. First, 200 ng DNA in 28 µL of $H_2O$ was used as input to the oxidation reaction, in which it was combined with 10 µL reconstituted TET2 reaction buffer + supplement, 1 µL oxidation supplement, 1 µL dithiothreitol (DTT), 1 µL oxidation enhancer and 4 µL TET2. This was mixed with 5 µL 0.4 mM Fe(II) solution and incubated for 1 h at 37 °C, after which 1 µL Stop reagent (ProtK) was added and incubated for 30 min at 37 °C. A 50 µL volume of oxidized DNA was cleaned with 90 µL AmpureXP beads (1.8x), denatured by boiling in formamide at 90 °C for 10 min, and quenched on ice. Deamination was performed by adding 10 µL APOBEC reaction buffer, 1 µL BSA and 1 µL APOBEC, bringing the volume to 100 µL with $H_2O$, and incubating at 37 °C for 3 h. A 100 µL volume of DNA was cleaned with 100 µL AmpureXP beads (1x) before library preparation.

Amplicon library preparation was performed in three rounds of PCR: amplification out of the converted DNA; addition of part of R1 and R2 sequencing primers; and addition of remaining sequencing handles and sample indexes. For each 50-µL PCR1 reaction, we used 20 µL converted DNA, 1.5 µL each 10 µM forward and reverse primers (oCR75 and oCR54), 2 µL H2O, and 25 µL RepliQa HiFi ToughMix (QuantBio, 95200-500) and ran the following thermocycling protocol: 30 s at 98 °C, then 22 cycles of 98 °C for 10 s, 56 °C for 30 s and 68 °C for 60 s, and then a final step of 68 °C for 5 min. PCR1 product was then cleaned with a 0.9x AmpureXP bead clean and eluted in 21 µL $H_2O$. For each 20-µL PCR2 reaction, we used 1 µL cleaned PCR1 product, 0.6 µL each 10 µM forward and reverse primers (oCR84 and oCR55) and 10 µL RepliQa HiFi ToughMix and ran the following thermocycling protocol: 30 s at 98 °C, then 10 cycles of 98 °C for 10 s, 55 °C for 5 s and 68 °C for 5 s, and then a final step of 68 °C for 1 min. PCR2 product was then cleaned with a 0.9x AmpureXP bead clean and eluted in 21 µL $H_2O$.

For each 50 µL PCR3 reaction, we used 22 µL cleaned PCR2 product, 1.5 µL each of 10 µM forward and reverse indexing primers, and 25 µL RepliQa HiFi ToughMix and ran the following thermocycling protocol: 30 s at 98 °C, then 6 cycles of 98 °C for 10 s and 68 °C for 5 s, and then a final step of 68 °C for 1 min. PCR3 product was then cleaned with a 0.9x AmpureXP bead clean and eluted in 17 µL $H_2O$. Libraries were then quantified using the Qubit 1X dsDNA HS Assay Kit for concentration and D1000 ScreenTape (Agilent Technologies, 5067–5582) for length. Libraries were pooled with ~15% PhiX Sequencing Control v3 (Illumina, FC-110-3001) for complexity and sequenced with 374 cycles on R1, 235 cycles on R2, 8 cycles on I1, and 8 cycles on I2 and with a MiSeq Reagent Kit v3 (600-cycle) (Illumina, MS-102-3003) on a MiSeq.

For amplicon-SMF analysis of the reporter gene, reads were aligned to a custom index built from the PCR amplicons using bwameth[5]. The resulting BAM files were filtered for alignment quality and uniqueness, unconverted reads were removed (by looking at the conversion of all non-GpC Cs), and bulk methylation was computed using MethylDackel (https://github.com/dpryan79/MethylDackel). Custom scripts were used to generate the single-molecule matrices and perform other quality control analyses. A Snakemake pipeline with associated conda environments is available at https://github.com/GreenleafLab/amplicon-smf. Molecules sharing identical methylation patterns across the entire amplicon were collapsed to a single representative, and all reported analyses use these deduplicated matrices.

Single-molecule states were assigned to each read using a maximum-likelihood approach, as previously described[2]. Briefly, all possible single-molecule states for a given amplicon were enumerated. Nucleosomes were mandated to start and end at GpCs, have at least one accessible GpC in a linker between any pair of nucleosomes, and be between 110 and 140 bp in length. For each hypothetical state, the expected methylation signal was computed by determining whether each GpC is accessible, protected by a nucleosome, or protected by a TF. These GpC occupancy states were then converted into probabilities using different methylation probabilities (unobserved) for each of the three possible occupancy states. This results in a matrix of size (number of states) × (number of GpCs) representing the probability that GpC $j$ is methylated in state $i$. This matrix is then converted into a matrix $P$ representing the probability of observing a converted base after the EM–seq reactions, estimated from bulk methylation measurements from MethylDackel. The maximum log-likelihood state for each input molecule, from single-molecule methylation matrix $M$, was computed using a Bernoulli likelihood across all GpC positions in a vectorized format ($\mathrm{argmax}(\log(P) \times M + \log(1 - P) \times (1 - M))$), resulting in a vector of the most likely state for each input molecule. To avoid introducing potential biases during preprocessing, we include all molecules in our downstream analyses, regardless of likelihood at the maximum-likelihood estimation. The code is available at: https://github.com/GreenleafLab/amplicon-smf/blob/master/workflow/scripts/classify_single_molecule_binding_v2.py.

To infer nucleosome occupancy from the deduplicated methylation matrices (Figure S7C), we quantified the average methylation frequency at GpC sites in between TetO sites for each condition.

*Clonal Cell Line Analysis*

To analyze the frequency of on-target reporter integration (Figure S1A) and transcriptional response functions of clonal cell lines (Figure S4I), we expanded a K-562 cell line containing a 5xTetO reporter gene (EC045) and expressing rTetR-VP48 (NVD060) and sorted single clones into separate wells of a 96-well tissue culture plate in 200 µL of RPMI media. We expanded each clonal line until confluent, then transferred 22 clones to a 24-well tissue culture plate and expanded until confluent. We then took a split of these cultures and added 0, 25, or 1000 ng/mL doxycycline to each line and measured gene expression over the next 5 days using flow cytometry as described above. One clone did not express the reporter gene and was discarded from all analysis.

At the start of the recruitment time course, we also took a split of cells from each line and extracted gDNA from ~1 million cells using the Monarch gDNA extraction kit. gDNA from a wild-type K-562 line was extracted in parallel. We then PCR-amplified the 5' AAVS1-reporter junction. For each reaction, we used 10 µL Q5 Hot Start High-Fidelity 2X Master Mix, 1 µL of each 10 µM forward and reverse primers (dna803 and dna804), and 30 ng of gDNA. PCRs were performed with the following thermocycling protocol: 30 s at 98 °C, then 30 cycles of 98 °C for 10 s, 70 °C for 30 s, and 72 °C for 30 s, and then a final step of 72 °C for 2 min. Amplified DNA was then visualized on an agarose gel made with SYBR Safe DNA Gel Stain (ThermoFisher, S33102).

*Estimating Citrine Degradation Rate*

To estimate Citrine degradation rates, we expressed a synTF with the strong KRAB repression domain from ZNF10 (JT151) in K-562 cells containing a genomically integrated 9xTetO reporter gene with the pEF promoter, which is constitutively active (JT039), as described above. This synTF has been shown to be an extremely rapid repressor in mammalian cells[6]. We then added 1000 ng/mL dox and measured gene expression dynamics over time using flow cytometry, as described above. To remove background silenced cells from downstream analysis, we set a gate next to this population and calculated the fraction of background silenced cells. From each subsequent day of the time course, we randomly removed this fraction of the population from cells with Citrine expression below this gate. We modeled Citrine degradation dynamics (from the mean Citrine fluorescence on each day) as a first order process with an offset to account for basal fluorescence from the cytometer using the equation $Citrine = Ae^{-kt} + C$. In this equation, $k$ is a combination of active degradation, $k_{d,prot}$, and effects from dilution through cell growth. We assumed a cell doubling time of 24 hours to calculate the active degradation rate. We fit this model to our recruitment time course data using the SciPy curve_fit function. Model parameters and fits are in Supplementary Table 2.

*Hill Function Fits to Transcriptional Response Functions*

For most analyses (Figures 2E-F, 4E) Hill functions of the form:

$$f(x) = \frac{x^n}{x^n + {EC_{50}}^n} \tag{1}$$

Were fit to transcriptional response functions using the SciPy curve_fit function. Here, $x$ is the independent variable (dox concentration or number of binding sites) and $f(x)$ is an aspect of the transcriptional response (either fraction of active cells or MFI of active cells). $n$ is the Hill exponent, which describes the apparent cooperativity, and $EC_{50}$ is the amount of input that produces a half-maximal response. For these analyses, transcriptional responses were normalized to fall between 0 and 1.

For average transcriptional responses following transient transfections in HEK-293T reporter cells, Hill functions of the form:

$$f(x) = \frac{Ax^n}{x^n + {EC_{50}}^n} + C \tag{2}$$

Were fit to transcriptional responses using the SciPy curve_fit function. In this case, data were not normalized and the parameter absolute_sigma was set to False. $x$ is the average mScarletI3 fluorescence for a particular bin, $n$ and

$EC_{50}$ are as described above, $A$ is the maximum value of the response, and C is the background fluorescence measured by the cytometer. Model parameters and fits are in Supplementary Table 2.

*Two-State Model of Gene Regulation*

Gene expression states in the two state bursting model shown in Figure 3A-C and Figure S3A can be described by the following system of differential equations:

$$\frac{dA}{dt} = k_A S - k_s A \tag{3}$$

$$\frac{dS}{dt} = -k_A S + k_s A \tag{4}$$

With the constraint $A + S = 1$. Here, $k_A$ is the rate of activation, $k_S$ is the rate of silencing, $A$ is the Active state, and $S$ is the Silent state. mRNA and protein levels can be described by the equations:

$$\frac{d[mRNA]}{dt} = Ak_{trx} - [mRNA]k_{d,rna} \tag{5}$$

$$\frac{d[Protein]}{dt} = [mRNA]k_{trl} - [Protein]k_{d,prot} \tag{6}$$

Here, $k_{trx}$ is the rate of transcription, $k_{trl}$ is the rate of translation, $k_{d,rna}$ is the mRNA degradation rate, and $k_{d,prot}$ is the protein degradation rate. To simulate transcriptional responses from these equations, we discretized the above equations for use with the Gillespie algorithm[7]. We included cell division explicitly in this model (such that $k_{d,prot}$ and $k_{d,rna}$ are the active degradation rates) by assuming that protein and mRNA levels are reduced by half every 24 hours. To un-synchronize cells, we assumed that the first division for each cell occurred at a random time between 0 and 24 hours. To reduce simulation run times, we pre-defined the number of reaction steps set to occur and pre-sampled random numbers for each reaction number and timestep. Simulations were stopped when the simulation timestep reached a specified value, and the number of reaction steps was always chosen to not be the limiting factor of the simulation. In order to better visualize bimodal protein expression distributions, we assumed that a leaky transcription rate of 0.5/hour occurred in the Silent state (otherwise, all Silent cells would have exactly 0 proteins expressed). Single-cell protein trajectories were obtained by saving the current protein level for a pre-defined number of snapshots evenly spaced across the simulation runtime. Model parameters are in Supplementary Table 2.

To estimate the rates of re-equilibration for sorted cell populations (Figure 3E; Figure S3D-E), we assumed that protein degradation was fast enough such that the fraction of cells in each protein expression state (Citrine On or Off) closely tracked the fraction of cells in the corresponding gene state (Active or Silent). This assumption would need to be relatively true in order to observe clearly separated populations of On and Off cells in the population, and allows us to only consider the equations describing $A$ and $S$ (above) when fitting our gene expression data. We also assumed that there is a characteristic lag time ($t_{lag}$) after dox addition until changes in the fraction of On cells is observed, which is related to the time it takes for enough protein to accumulate for a cell to be called as On from our cytometry gate. In this case, and assuming that all cells start in the S state, the system of equations has the analytic solution:

$$A(t) = \frac{k_A}{k_A + k_S}\left(1 - e^{-(k_A + k_S) \times max(t - t_{lag}, 0)}\right) \tag{7}$$

We fit this solution to the pre-sort fraction of On cells (Figure 3E) using the SciPy curve_fit function. Model parameters and fits are in Supplementary Table 2. To predict expected gene expression dynamics from sorted populations, we used these fit parameters to solve the system of differential equations with the initial conditions A = 1 and A = 0 for sorted On and sorted Off populations respectively. Model parameters and fits are in Supplementary Table 2.

*Locus-Level vs. Cell-Level Variability Decomposition*

To estimate the relative contribution of locus-level and cell-level effects on variability in transcriptional response functions, we closely followed Elowitz and colleagues[1] but for binary variables (Off vs. On reporters). We used the expressions:

$${\eta_{locus}}^2 = \frac{P(c \neq r)}{2P(c)P(r)} \tag{8}$$

$${\eta_{cell}}^2 = \frac{P(c \,\&\, r) - P(c)P(r)}{P(c)P(r)} \tag{9}$$

Where $\eta$ is variability and *c* and *r* are binarized expression states for the Citrine and mCherry reporter respectively (On = 1, Off = 0). *P* indicates probabilities, i.e. *P*(*c*) is the probability of a cell expressing Citrine. To calculate the fraction of variability from each source, we assumed reporters activated symmetrically and thus ${\eta_{total}}^2 = {\eta_{locus}}^2 + {\eta_{cell}}^2$.

*Combined Kinetic and Thermodynamic Model of Gene Regulation*

We sought to develop a mathematical model of gene regulation that connected microscopic features, such as synTF binding, to macroscopic changes in gene expression in order to link locus-level variability to population-level transcriptional response functions in the Tet-On system. Thermodynamic models are powerful tools for describing gene regulation while explicitly considering variables such as TF occupancy[8,9]. These models assume that the system of interest can exist in a finite set of discrete microstates. If the system is in thermodynamic equilibrium, statistical mechanics can be used to calculate the probability of the system occupying any one microstate based on the relative energies of each state (which is determined by molecule concentrations, binding affinities, and other features of the system). We developed a combined kinetic and thermodynamic model of gene regulation where synTF occupancy was used as inputs to the rates $k_A$ and $k_{trx}$ in the kinetic framework described by Equations 3 to 6.

We considered a coarse-grained approximation of a reporter gene with 9 TetO sites, which can be bound by synTFs, and a proximal promoter site where RNA polymerase (or more abstractly, general transcriptional machinery) could bind. Based on the recently proposed "nucleosome destabilization" model[2], we assumed that synTFs compete with nucleosomes for binding at TetO sites, and once bound destabilize nearby nucleosomes. We also assumed that a nucleosome could occupy the promoter site, and for simplicity that a single nucleosome covers 3 adjacent TetO sites when bound (except for edge cases at the ends of the TetO arrays).

To determine average synTF occupancy, it is useful to calculate the partition function of the system, which is obtained by summing the statistical weights of every possible microstate (i.e., binding configuration of synTFs, nucleosomes, and polymerase). Therefore, we recursively enumerated every possible binding state and tracked the number of synTFs bound, number of nucleosomes bound, and whether or not polymerase was bound for each state. Then, we calculated the respective weight of each state using the equation:

$$W = \left(\frac{[TF]}{K_{TF}}\right)^{N_{TF}} \left(\frac{[Nuc]}{K_{Nuc}}\eta^{H(N_{TF})}\right)^{N_{Nuc}} \left(\frac{[Pol]}{K_{Pol}}\right)^{N_{Pol}} (\lambda^{N_{TF}N_{Pol}}) \tag{10}$$

$$W = \left(\frac{[TF]}{K_{TF}}\right)^{N_{TF}} \left(\omega_{Nuc}\eta^{H(N_{TF})}\right)^{N_{Nuc}} (\omega_{Pol})^{N_{Pol}} (\lambda^{N_{TF}N_{Pol}}) \tag{11}$$

Here, $[TF]$ is the abundance of the synTF (which we assume to be directly proportional to dox concentration and work in units of ng/mL), $[Nuc]$ is the abundance of nucleosomes, $[Pol]$ is the abundance of polymerase, and $K_{TF}$, $K_{Nuc}$, and $K_{Pol}$ are their respective dissociation constants. $N_{TF}$, $N_{Nuc}$, and $N_{Pol}$ are the number of each molecule bound in a particular microstate between zero and the maximum possible at this promoter (1 for polymerase, 9 for synTFs, and 5 for nucleosomes). In Equation 11, we have substituted in the dimensionless parameters $\omega_{Nuc} = [Nuc]/K_{Nuc}$ and $\omega_{Pol} = [Pol]/K_{Pol}$ for some of these variables. To represent the effect of active nucleosome remodeling by the synTF, as observed previously[2], we introduce the parameter $\eta < 1$ which modifies the binding weight of nucleosomes if a synTF is bound at the reporter ($H(N_{TF})$ is the Heaviside function, i.e., 0 if there are no synTFs bound and 1 if there is at least one bound). $\lambda > 1$ represents a favorable interaction between synTFs and polymerase when both are bound at the reporter, representing the regulatory effect of the synTF.

The partition function of the system is calculated by summing the statistical weights of every microstate:

$$Z = \Sigma_s W_s \tag{12}$$

From the partition function, the probability of *TF* synTFs being bound is:

$$P(TF) = \frac{1}{Z}\Sigma W_{N_{TF} = TF} \tag{13}$$

Where $\Sigma W_{N_{TF} = TF}$ is the total statistical weight of states with *TF* synTFs bound. Average synTF occupancy is then:

$$\langle TF \rangle = \sum_{TF = 0}^{N} P(TF)TF \tag{14}$$

Where *N* is the number of binding sites in the reporter gene. Similarly, the average polymerase occupancy is:

$$\langle Pol \rangle = \frac{1}{Z}\Sigma W_{N_{Pol} = 1} \tag{15}$$

As commonly done with thermodynamic models, we assumed that there was a separation of timescales between TFs binding and unbinding and transcription, and therefore that the rate of transcription ($k_{trx}$) was proportional to the average occupancy of RNA polymerase[10]:

$$k_{trx} = \beta \ \langle Pol \rangle \tag{16}$$

Where $\beta$ is a proportionality constant. However, from our single-molecule footprinting data, we know that gene activation rate does not have the same relationship with synTF occupancy as the rate of transcription, which saturates at a much higher number of synTFs bound (Figure 6H). Therefore, we assume that the rate of activation is related to average synTF occupancy through a Hill function:

$$k_A = \frac{\alpha \ \langle TF \rangle^{\ n}}{EC_{50}{}^{n} \ + \ \langle TF \rangle^{\ n}} \tag{17}$$

Where $\alpha$ is a proportionality constant, $n$ is the Hill exponent, and $EC_{50}$ is the average number of synTFs bound where $k_A$ is half-maximal.

By combining Equations 16 and 17 with Equations 3 to 6, we obtained a kinetic model of gene regulation where the rates of activation and transcription were obtained from the microscopic binding states of the reporter gene, which in turn were determined by the primary experimental variable $[TF]$, which we assumed was proportional to the dox concentration.

<u>*Fitting Kinetic and Thermodynamic Model to Population Responses*</u>

Given the combined kinetic and thermodynamic model of transcriptional regulation in the Tet-On system described above, we wondered if incorporating variability into key parameters governing the transcriptional responses could reproduce both single-cell and population-level experimental observations. From our experimental results, we observed that variability in transcriptional response functions are encoded at the locus-level and associated with changes in synTF binding and activation strength when bound. We therefore assumed that the nucleosome binding parameter $\omega_{Nuc}$ and the synTF binding threshold at which the gene activates, $EC_{50}$, varied between cells. Variability in $\omega_{Nuc}$ represents the effect of the local chromatin environment on synTF binding, and variability in the $EC_{50}$ represents the effect of the local chromatin environment on the activation strength of the synTF when bound.

Because cells with less activation potential had both reduced synTF binding and activation strength when bound, we assumed that the values of these parameters in the cell population followed a multivariate log-normal distribution:

$$\big(ln(\omega_{Nuc}), ln(EC_{50})\big) \sim \mathcal{N}(\mu \,, \Sigma) \tag{18}$$

$$\mu \ = \frac{ln\langle \omega_{Nuc} \rangle \ - \ \frac{1}{2} ln\left(1 + \frac{\sigma_{Nuc}^2}{\langle \omega_{Nuc} \rangle^2}\right)}{ln\langle EC_{50} \rangle \ - \ \frac{1}{2} ln\left(1 + \frac{\sigma_{EC50}^2}{\langle EC_{50} \rangle^2}\right)} \tag{19}$$

$$\Sigma \ = \frac{ln\left(1 + \frac{\sigma_{Nuc}^2}{\langle \omega_{Nuc} \rangle^2}\right), \quad \rho \sqrt{ln\left(1 + \frac{\sigma_{Nuc}^2}{\langle \omega_{Nuc} \rangle^2}\right) ln\left(1 + \frac{\sigma_{EC50}^2}{\langle EC_{50} \rangle^2}\right)}}{\rho \sqrt{ln\left(1 + \frac{\sigma_{Nuc}^2}{\langle \omega_{Nuc} \rangle^2}\right) ln\left(1 + \frac{\sigma_{EC50}^2}{\langle EC_{50} \rangle^2}\right)}, \quad ln\left(1 + \frac{\sigma_{EC50}^2}{\langle EC_{50} \rangle^2}\right)} \tag{20}$$

Where $\langle \omega_{Nuc} \rangle$ and $\langle EC_{50} \rangle$ are the linear-space means of each variable, $\sigma_{Nuc}$ and $\sigma_{EC50}$ are the linear-space standard deviations, and $\rho$ is the log-space correlation coefficient between the two variables.

We then fitted predicted transcriptional responses from this stochastic model to the experimentally observed fraction of active cells and mean expression level of active cells. To do this, we randomly sampled 300 pairs of $\omega_{Nuc}$ and $EC_{50}$ from Equations 18 to 20 for a set of $\langle \omega_{Nuc} \rangle$, $\langle EC_{50} \rangle$, $\sigma_{Nuc}$, and $\sigma_{EC50}$ values (which were free parameters in the fit). We assumed $\omega_{Nuc}$ and $EC_{50}$ values were correlated with a correlation coefficient of $\rho = 0.6$ because cells with lower activation potentials had both reduced synTF binding and weaker activation strength when bound (Figure 6G-H).

For each pair of $\omega_{Nuc}$ and $EC_{50}$ values, we then calculated $k_A$ and $k_{trx}$ according to Equations 16 and 17 for a range of synTF input levels that matched the dox concentrations in our experimental data and assuming that $K_{TF}$ = 20 ng/mL dox and $\eta = 0.8$, which we estimated empirically by comparing predicted binding distributions with our SMF data (Equation 11). The free parameters in this step were $\omega_{Pol}$, $\lambda$, and $n$ (the Hill Exponent in Equation 17). We normalized our experimental data so that each aspect of the transcriptional response varied between 0 and 1, allowing us to set $\alpha = 1$ and $\beta = 1$ in Equations 16 and 17.

To calculate the fraction of active cells and their mean expression level from the predicted set of $k_A$ and $k_{trx}$ values at each dox concentration, we computed the soft threshold weight:

$$w_i = \frac{1}{1 + e^{-45(k_A - 0.9)}} \tag{21}$$

For each $k_A$ value. This is a switch-like, but smooth, function that sharply increased from 0 to 1 at the threshold $k_A$ = 0.9 was necessary to implement so that we could effectively fit the model to our data using sciPy curve_fit. We then computed:

$$Frac\ On = \frac{1}{N\ cells} \sum_{i=1}^{cells} w_i \tag{22}$$

Where *i* denotes each cell in the population, to obtain a proxy for the fraction of active cells in the population. This approximation is accurate when $k_A$ is much greater than $k_S$ (Equations 3 to 4), which we obtain by setting $k_S$ = 0.1/hour and $\alpha = 40$/hour for our stochastic simulations.

Finally, we computed the mean transcription rate of active cells by weighting the predicted $k_{trx}$by equation 21:

$$On\ Cell\ k_{trx} = \sum_{i=1}^{cells} w_i k_{trx,i} \tag{23}$$

We normalized the predicted On cell transcription rates to have a maximum of 1. Thus, for each dox concentration we obtained the predicted fraction of active cells and predicted mean expression level of active cells for a certain parameter set $\{\langle \omega_{Nuc} \rangle, \langle EC_{50} \rangle, \sigma_{Nuc}, \sigma_{EC50}, \omega_{Pol}, \lambda, n\}$ and could compare these predicted values to our experimental observations (Figure 2E). We then optimized these predictions using the SciPy curve_fit function.

To obtain predicted synTF binding distributions for sorted On and Off cells (Figure 7E), we randomly sampled 1000 pairs of $\omega_{Nuc}$ and $EC_{50}$ values using the optimal set of parameters obtained by fitting, $\{\langle \omega_{Nuc} \rangle, \langle EC_{50} \rangle, \sigma_{Nuc}, \sigma_{EC50}, \omega_{Pol}, \lambda, n\}$ and $[TF]$ = 14 ng/mL (which resulted in 50% cells On), then calculated corresponding

$k_A$ and $k_{trx}$ values as described above. We then separated cells (i.e., pairs of $k_A$ and $k_{trx}$ values) into On or Off conditions based on the threshold $k_A$ > 0.9. For each cell, we calculated the predicted synTF binding distribution from Equation 13, then summed and normalized these distributions for On and Off cells to generate the ensemble predicted synTF binding distribution for each respective cell population.

<u>*Simulating Gene Expression Dynamics from Combined Model*</u>

To simulate gene expression dynamics from our combined kinetic and thermodynamic model, we randomly sampled 500 pairs of $\omega_{Nuc}$ and $EC_{50}$ values using the optimal set of parameters obtained by fitting, $\{\langle\omega_{Nuc}\rangle$, $\langle EC_{50}\rangle$, $\sigma_{Nuc}$, $\sigma_{EC50}$, $\omega_{Pol}$, $\lambda$, $n\}$, then calculated corresponding $k_A$ and $k_{trx}$ values as described above. We discretized Equations 3 to 6 and simulated gene expression dynamics using the Gillespie algorithm[7] as described above (including cell division and pre-sampling random numbers) then simulated gene expression dynamics for each cell. To better visualize the Off population of cells, we assumed a leaky transcription rate of 1/hour in the Silent state. Full model parameters can be found in Supplementary Table 2.

For *in silico* cell sorting and transient dox upshifts, we simulated gene expression dynamics as described with [*TF*] = 35 ng/mL dox. After 300 hours of simulation time, [*TF*] was increased to 1000 ng/mL, then reverted to 35 ng/mL at the 400 hour mark. After running the simulation for all cells, protein trajectories were analyzed at the timepoint closest to 100 hours for cell sorting. Cells with a protein level at this time greater than 800 were designated as On, cells with a protein level less than 300 were designated as Off. The gene expression dynamics of sorted cell populations were then compared.
